# The SPICA Coarse-Grained Force Field for Proteins and Peptides

**DOI:** 10.1101/2021.12.01.470839

**Authors:** Shuhei Kawamoto, Huihui Liu, Sangjae Seo, Yusuke Miyazaki, Mayank Dixit, Russell DeVane, Christopher MacDermaid, Giacomo Fiorin, Michael L. Klein, Wataru Shinoda

## Abstract

A coarse-grained (CG) model for peptides and proteins was developed as an extension of the SPICA (Surface Property fItting Coarse grAined) force field (FF). The model was designed to examine membrane proteins that are fully compatible with the lipid membranes of the SPICA FF. A preliminary version of this protein model was created using thermodynamic properties, including the surface tension and density in the SPICA (formerly called SDK) FF. In this study, we improved the CG protein model to facilitate molecular dynamics (MD) simulation with a reproduction of multiple properties from both experiments and all-atom (AA) simulations. The side chain analogs reproduced the transfer free energy profiles across the lipid membrane and demonstrated reasonable dimerization free energies in water compared to those from AA-MD. A series of peptides/proteins adsorbed or penetrated into the membrane simulated by the CG-MD correctly predicted the penetration depths and tilt angles of peripheral and transmembrane peptides/proteins comparable to those in the orientation of protein in membrane (OPM) database. In addition, the dimerization free energies of several transmembrane helices within a lipid bilayer were comparable to those from experimental estimation. Application studies on a series of membrane protein assemblies, scramblases, and poliovirus capsids demonstrated a good performance of the SPICA FF.

## 1. INTRODUCTION

Molecular dynamics (MD) simulation is a powerful tool for investigating the structure and dynamics of biomolecular complexes, including lipids and proteins. All-atom (AA) MD simulations with explicit solvents and ions provide useful information for predicting various chemical properties, such as the partitioning of small molecules. However, even with the fastest currently available computers, atomistic simulations remain several magnitudes away from being able to effectively cover the spatial and temporal scales of most membrane protein processes. To address this challenge, coarse grained (CG) models have been proposed and used. In CG models, several atoms are grouped into a single particle, and the interactions between these CG particles are described by effective potentials that incorporate both energetic and entropic contributions resulting from integrating out (or averaging over) the neglected atomistic details. Coarse-graining greatly enhances computational efficiency for exploring the reduced phase space. For example, mapping from three or four heavy atoms into a single CG particle using a top-down approach (using the prepared, simplified functions for intermolecular interaction) accelerates MD simulations by three orders of magnitude.^1^ The MD simulation of the CG model is a potential method by which to investigate processes involving multiple membrane proteins.

Many CG protein models have been developed with various resolutions for aims such as structure prediction, protein folding and aggregation, protein-protein recognition and docking and the simulation of various proteins.^2, 3, 4–11, 12, 13^ The CG modeling of membrane proteins is relatively less studied. To further decrease the number of CG particles in the systems, implicit solvent and/or lipid models have been used for the majority of CG models of membrane proteins.^14, 15, 16–23, 24, 25^ MARTINI is the most popular CG model with explicit water and lipids. It uses one bead for protein backbone and a 4:1 mapping for protein sidechains and lipids.^26–28^ PACE is a finer protein model which can be incorporated with MARTINI water and lipids models.^29–32^ The atomistic representation of backbone and united atoms in the sidechains can simulate the formation of hydrogen bonds.^29^ SIRAH is another CG protein model with explicit WT4 water modeling. In the SIRAH model, backbones are represented by three particles and sidechains by up to five particles.^33^ A compatible CG phospholipids model was introduced for SIRAH to explore membrane-protein dynamics.^34^ Even though these CG models are currently available, we were motivated to develop a CG protein model in compatible with Surface Property fItting Coarse grAined (SPICA) water and lipid force field (FF) (formerly known as the SDK models),^1, 35, 36^ due to the advantageous features of SPICA lipid membranes.

The SPICA FF provides an accurate description of lipid membrane properties, based on a systematic CG parametrization to reproduce surface/interfacial tension, density, and solvation/transfer free energy as well as structure, namely, distribution functions obtained from all-atom MD simulations.^1, 35, 36^ Owing to the reproduction of the elastic properties and line tension of the membranes, the SPICA model provides a promising way to investigate the mesoscopic morphological variation of lipid membranes.^37–39^ The development of the SPICA(SDK) CG protein model was initiated by some of the present authors to predict native structures from protein decoy data sets.^40^ In this preliminary version, amino acids were represented by single backbone particles and 0-3 side chain particles and the non-bonded parameters were optimized by reproducing the surface tension and density of side chain analogs. Thus, the partial CG model can be expected to be transferable. However, the CG model require further efforts to complete the bond parameters for backbones and the non-bonded parameters between proteins and lipids.

In this study, we improved and complemented the preliminary SPICA(SDK) protein model by reproducing multiple properties from both experiments and MD simulations. To better reproduce the structure of the protein, the side chains of PHE, TYR were remapped to four CG particles and HIS to 3. Bond parameters were taken from reference structures from the PDB database or equilibrated AA MD. The initial secondary structures were maintained during the simulations by employing an elastic network model.^41^ To reproduce the interaction between protein and lipid, non-bonded side chain parameters were optimized to reproduce the hydration free energy, dimerization free energy of side chain analogs in water, and the partitioning free energy of side chain analogs between water and dioleoyl phosphatidylcholine (DOPC) membrane. Subsequently, the non-bonded parameters of the backbones were optimized to reproduce the penetrating depth and tilt angle of peripheral peptides/proteins. Lastly, to improve the effective interaction between proteins, the dimerization free energies of the transmembrane helices were calculated to optimize the backbone parameters. The SPICA protein model succeeded in predicting native structures from decoy sets, simulating the assembly of membrane proteins in the DOPC lipid bilayer. Several application studies demonstrated that the SPICA FF is useful for MD simulation of proteins in different environments.

## 2. METHODS

### 2.1 Coarse-grained Force Field

The SPICA protein model is an extension of the previous amino acid model proposed by DeVane et al.^40^, where the parameter was optimized based on thermodynamic data including density and surface tension. The model is fully compatible with the lipid and solvent models within the SPICA FF.^1, 35^ Here we further optimized the interacting parameter set using a slightly different definition of the CG segments.

The CG mapping for all naturally occurring amino acids is shown in Figure 1. Briefly, the backbone (BB) atoms of one amino acid are represented by one CG particle centered at its alpha-carbon atom, while the side chain (SC) is composed of 0-4 CG particles centered at the center of mass of the corresponding heavy atoms, according to their size and shape. The major differences from the previous model^40^ are as follows: (i) a four-beads representation was incorporated for the phenyl-based SC of PHE and TYR to better simulate the planar structure; (ii) the HIS SC was parameterized using three CG particles.

**Figure 1.**
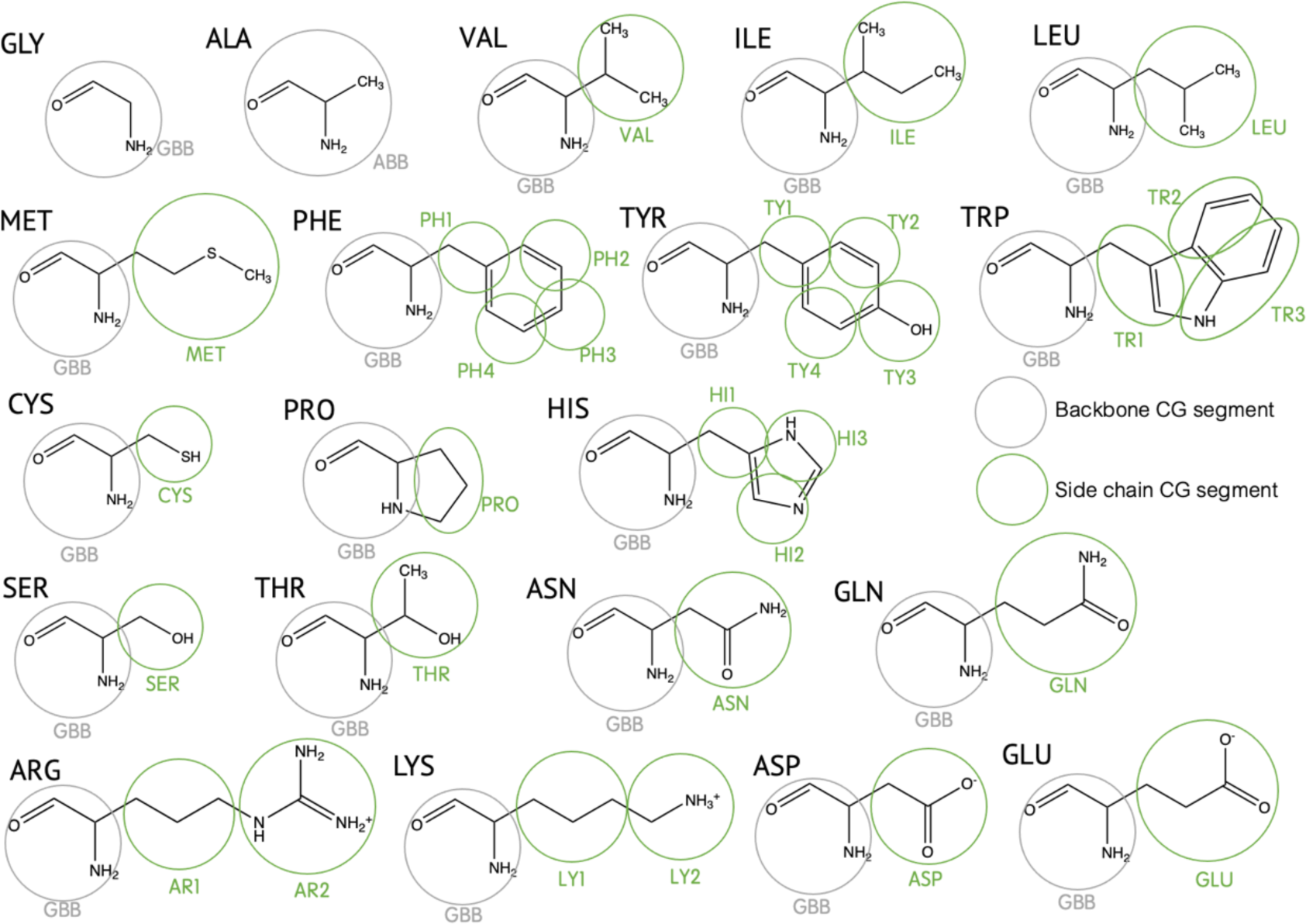
CG mapping of protein for 20 amino acids in the SPICA FF. Segment names are shown in gray for backbone and green for side chain segments.

In this study, the SPICA protein model was designed to investigate the properties of membrane proteins, such as the assembly of transmembrane peptides and the deformation of the lipid bilayer by membrane proteins. However, it can also be used for capsid proteins, as shown later. Therefore, the CG protein model should retain the correct structure, position, orientation, and association energy of a membrane protein in the lipid bilayer. The first property, the structure of a protein is mainly related to bond interactions, while the others are dominated by non-bonded interactions.

The bond interactions are described by simple harmonic potentials.^1^ The bond stretching, angle bending and dihedral potentials are given by Eqs. (1)-(3),

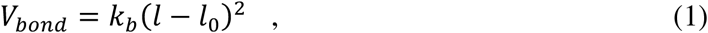

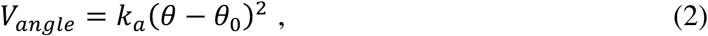

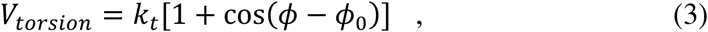

where *k_b_*, *k_a_*, and *k_t_* are the force constants, and *l*_0_ and θ_0_ are the energy minimum values for the bond length and angle, respectively. ϕ_0_ is the phase of the dihedral angle. The angle potential is used for every angle created by connecting two bonds, except for the bonds in triangular SCs. The torsion potential is only used for SCs of PHE and TYR to maintain planar structures (Figure S1).

The SPICA FF employs simple Lennard-Jones (LJ)-type non-bonded potential functions. To reproduce the structural and interfacial properties of water/alkane systems, from our previous experience, LJ (12-4) functional form was likely to be the best choice for the pairs involving water, and LJ (9-6) for any others.^1^ In which all pairs involving protein particles interact via the LJ (9-6) potential described in Eq. (4), mainly for ease of implementation,

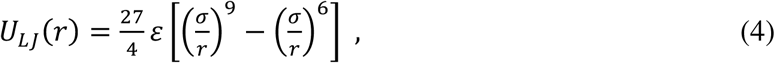

where *ε* is the minimum energy and σ is the distance when *U* = 0. The LJ potential for 1-2 and 1-3 bonded pairs were excluded from the calculation. A simple truncation was used for the long-range LJ interactions without any smoothing or shifting. A long cutoff distance of 15 Å was chosen to prevent significant artifacts from this truncation.

Electrostatic interactions are considered between ionic CG segments, which are found in charged residues (ARG, LYS, ASP and GLU) and the terminal groups of proteins. The interaction between charged particles *i* and *j* is given as

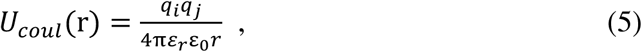

where ε_0_ is the permittivity in vacuum. The relative permittivity, ε_*r*_, was chosen to be 80 in the SPICA FF.^42^ To reproduce the correct ionic distribution, a non-cutoff scheme, such as PPPM,^43^ is used for Columbic interactions.

Considering our aim of reproducing the correct protein structures in simulations, we employed an elastic network model (ENM) to maintain the secondary structures. A harmonic bond is applied on two BB particles when their spatial distance is less than 9 Å, or they are separated by more than two bonds. As suggested in a previous study,^44^ a harmonic spring constant of 1.195 kcal/mol/Å^2^ was used. The cutoff distance and spring constant for the ENM may be changed depending on the target system; however, in this study, we continued to use the same values.

## 2.2 Parameter Optimization

The parameters for the bond stretching potential were determined by analyzing the bond distributions of 500 target proteins from the PDB database, except for the parameters for residues PHE, TYR, and TRP, which were taken from a previous paper.^40, 45^ The average bond length from the bond distribution was used as the equilibrium length *l*_0_. The force constant was obtained using the equation 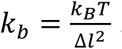, where *k_B_* is the Boltzmann constant, *T* is the temperature, and Δ*l* is the standard deviation of the bond length distribution. *k_b_*was limited to a maximum value of 50 kcal/mol/Å^2^ for the numerical stability of the MD simulation.

The parameters for the angle bending potential were obtained in a similar manner. However, the parameters for angles within the SCs of PHE and TYR were taken from a previous study.^45^ The force constant was calculated by using the equation 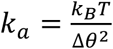, where Δ*θ* is the standard deviation of the angle distribution. To reproduce the correct protein structures with various secondary structures, the equilibrium angle θ_0_ for angles involving BB particles was set to the value calculated from the corresponding starting structure (e.g. PDB structure).

The torsion potential was considered only in the SCs of the PHE and TYR. The force constant *k*_*t*_ was set to 50 kcal/mol, while the equilibrium dihedral angle ϕ_0_ was calculated from the initial structure.

The LJ non-bonded parameters for proteins were optimized for multiple properties evaluated by either experiment or AA MD (Table 1 and 2). In a previous work, the LJ parameters for both SCs and BBs were fixed with surface tension and density and the cross interactions employed the Lorentz-Berthelot combination rules (geometric average for epsilon and arithmetic average for sigma).^40^ However, the preliminary model could not reproduce several thermodynamic quantities, such as the solvation free energies of SC analogs. Therefore, we decided to further optimize the LJ *ε*-parameters systematically using the protocol summarized in Table 1 and 2.

**Table 1.** Target properties or parameterization rules for LJ parameters.

| Scheme # | Target properties or parameterization rules |
| --- | --- |
| I | Surface tension and density of side chain analog liquid<sup>40</sup> |
| II | Combination rule (cross term for $\sigma$; arithmetic average) |
| III | Hydration free energy of side chain analog |
| IV | Hydration free energy of peptides |
| V | Radial distribution function of side chain dimers in water |
| VI | Free energy profile of side chain analog across lipid bilayer |
| VII | Penetration depths and tilt angles of peripheral peptides/proteins |
| VIII | Tilt angle of transmembrane peptide WALP |
| IX | Dimerization free energy of transmembrane helices |

First, the interaction between water and protein was optimized by reproducing the hydration free energy (Scheme III in Table 2). For SCs, the experimental hydration free energies of the SC analogs were used as a reference. Since there is no available experimental data for BBs, we chose the hydration free energies of the dipeptide and polypeptide helix (8 residues) from AA MD as an alternative.

**Table 2.** Parameterization of LJ potential for pairs of particle types. The numbers in the table refer to the scheme indexes in Table 1.

| | $\epsilon$ | | $\sigma$ | |
| --- | --- | --- | --- | --- |
|  | SC | BB | SC | BB |
| Side chain (SC) | I, V |  | I, V |  |
| Backbone (BB) | IX | IX | II | I |
| Water | III | VII, VIII | II | IV |
| Alkane (lipid tail) | VI | VII, VIII | II | II |
| Lipid head | VI | VII, VIII | II | II |

Order of optimization I, II → III, IV → V → VI, VII, VIII → IX

We then evaluated the dimerization free energies of pairs of SC analogs in water by comparing CG and AA MD. Even though we were able to obtain good results for almost all pairs, this does not provide a good reference for the effective interaction between protein SCs, especially for heterogeneous molecules, such as ARG and LYS. This is because a strong association between two heterogeneous SC analogs take place when facing the hydrophobic segments with each other, though this is not allowed in the protein SCs because the hydrophobic segments typically connect to the protein backbones. Then, we used the connecting dimer of two amino acid SC analogs. To simplify the computation, we carried out AA-MD of 24 SC dimers in water (635-667 molecules) and used the calculated radial distribution functions (RDF) as a reference for the CG parameter optimization between SC segments. (Scheme V)

Next, the interactions between proteins and lipids were optimized. The LJ parameters between SCs and lipid were optimized to reproduce the free energy profiles of SC analogs along the bilayer normal in AA MD. This ensures that the SCs have correct partitioning between the hydrophilic and hydrophobic environments. After the optimization of SCs, the LJ parameters between the BBs and lipid segments were optimized to reproduce the correct position and orientation of peptides/proteins on/in the lipid bilayer. Specifically, the penetration depths and tilt angles of the peripheral alpha-helical peptides and proteins from the orientations of proteins in membranes (OPM) database,^46^ as well as the tilt angle of transmembrane peptide WALP from both experiments and AA MD were used as a reference. The penetration depths of peptides/proteins in the lipid bilayer are determined by the atom position reaching the deepest of the bilayer. The zero point of depth is the hydrophobic boundary of the bilayer, as in the OPM database, which is approximately 5 Å closer to the bilayer center than the position of the lipid phosphates (Figure S2). The tilt angle is defined as the angle between the long axis of the peptides/proteins and the bilayer normal (Figure S2). Considering the shielding effect of SCs on BBs, the LJ epsilons between the BBs and water were also adjusted in this step (Scheme VII).

Finally, the interaction parameters of the BB-BB and BB-SC segments were optimized to reproduce the dimerization free energies of transmembrane peptides, including glycophorin A (GpA), serine zipper (SerZip), WALP peptides, and the transmembrane segments of EphA1 and ErbB1. This parameterization process ensures reasonable associations of proteins in the lipid bilayer in the CG MD.

All of the parameters are available from the SPICA website (https://www.spica-ff.org/).

### 2.3 MD simulations

CG MD simulations were performed using the LAMMPS code.^47^ The initial CG protein structures were mapped from either crystal structures or equilibrated structures from AA MD using a converting tool, which is available on the SPICA website (https://www.spica-ff.org/). The energy minimization of the systems was conducted using the conjugate gradient algorithm, and the production simulations were run with a time step of 10 fs in the NPT ensemble. The pressure was maintained at 1 atm using the Parrinello-Rahman barostat,^48, 49^ while the temperature was set to 310 K (unless otherwise stated) by the Nosé-Hoover thermostat.^50, 51^ Notably, the temperature used was 303 K in the parameterization of our previous CG lipid, water and protein models.^1, 35, 40^ Strictly speaking, the CG models do not have transferability for temperature because the entropy of the CG model is different from that of the AA model.^52^ However, the discrepancy of parameters between 303 K and 310 K can be negligible in the optimization of the CG protein model.

The AA MD simulations were performed using GROMACS (ver. 5.1.4)^53^. The CHARMM36 force field^54, 55^ was employed in AA-MD to generate the reference trajectory data for training the CG parameters. The systems were energy-minimized using the steepest descent algorithm, and the production runs were carried out in the NPT ensemble. The pressure and temperature were the same as those for the corresponding CG system. Long-range electrostatic interactions were calculated using the Particle Mesh Ewald (PME) method.^56, 57^ LJ interactions were switched off by applying a force-switching function from 10 to 12 Å. All bonds, including hydrogens, were constrained by the LINCS algorithm^58^ to facilitate the use of a 2-fs timestep.

## 3. RESULTS AND DISCUSSION

The non-bonded LJ parameters were optimized to reproduce the target properties listed in Table 1. In this section, we discuss the performance of the optimized CG FF with respect to the target properties, either from experiments or AA MD.

### 3.1 Hydration free energies of side chain analogs

The hydration free energies of the SC analogs were calculated to optimize the LJ parameters between SCs and water. The analog molecules used for the side chains are listed in Table 3. Notably, VAL and PRO share the same analog, propane. The hydration free energies were calculated by pulling the side chains from vacuum to bulk water (1,000 molecules) along the normal to the vacuum-water interface. The PMF was computed using umbrella sampling^59^ and weighted histogram analysis method (WHAM).^60^ Twenty-five windows for a 45 Å distance were made, and each window was run for 10 ns. A simulation temperature of 298 K was selected to maintain consistency with the experimental conditions. The calculated hydration free energies using the SPICA CG model were in good agreement with the experimental values,^61^ as shown in Table 3. GLN exhibited the largest difference, which was still less than 0.5 kcal/mol.

**Table 3.** Hydration free energy, *ΔG*_hyd_, of side chains

| Residue | Side chain analog | $\Delta G_{\text{hyd}}$ (kcal/mol) | | Residue | Side chain analog | $\Delta G_{\text{hyd}}$ (kcal/mol) | |
| --- | --- | --- | --- | --- | --- | --- | --- |
|  |  | Exp. | SPICA |  |  | Exp. | SPICA |
| LEU | Isobutane | 2.28 | 2.5 | THR | Ethanol | -4.88 | -4.9 |
| ILE | Butane | 2.15 | 2.3 | SER | Methanol | -5.06 | -5.3 |
| VAL | Propane | 1.99 | 2.2 | TRP | 3-methylindole | -5.88 | -6.0 |
| PHE | Toluene | -0.76 | -0.6 | TYR | Cresol | -6.11 | -6.1 |
| CYS | Methanethiol | -1.24 | -1.1 | GLN | Propionamide | -9.38 | -8.9 |
| MET | Methyl-ethyl-sulfide | -1.48 | -1.5 | ASN | Acetamide | -9.68 | -9.9 |

The hydration free energies for the charged side chains (ARG, LYS, ASP, and GLU) were calculated using AA MD as a reference for parameterization (data not shown). However, in this approach, we needed to choose a large LJ epsilon value to observe the CG water crystallization around the charged SCs to reproduce high hydration free energies. This problem was difficult to solve as long as we used a non-polar CG water model. Therefore, for the charged SCs-water interaction of the present CG FF, we chose to use the RDF of SC dimers in water from AA MD to optimize the interactions between charged SCs and water, as will be presented in the following section.

### 3.2 Interactions between side chains in water

Starting with the preliminary set of LJ parameters between SC segments employing the combination rules, we optimized the LJ parameters to reproduce the dimerization free energies of two SC analogs in water from AA MD for most pairs. As shown in Figure 2, the PMFs of dimerization from AA (black lines) and CG (red lines) MD almost overlapped for pairs of polar, hydrophobic, and ring side chains. For charged and HIS side chains, consistent PMFs were also observed for some pairs, such as ARG-PRO, ARG-THR, and ARG-VAL, while the other pairs showed large deviations from AA MD (Figure S3).

**Figure 2.**
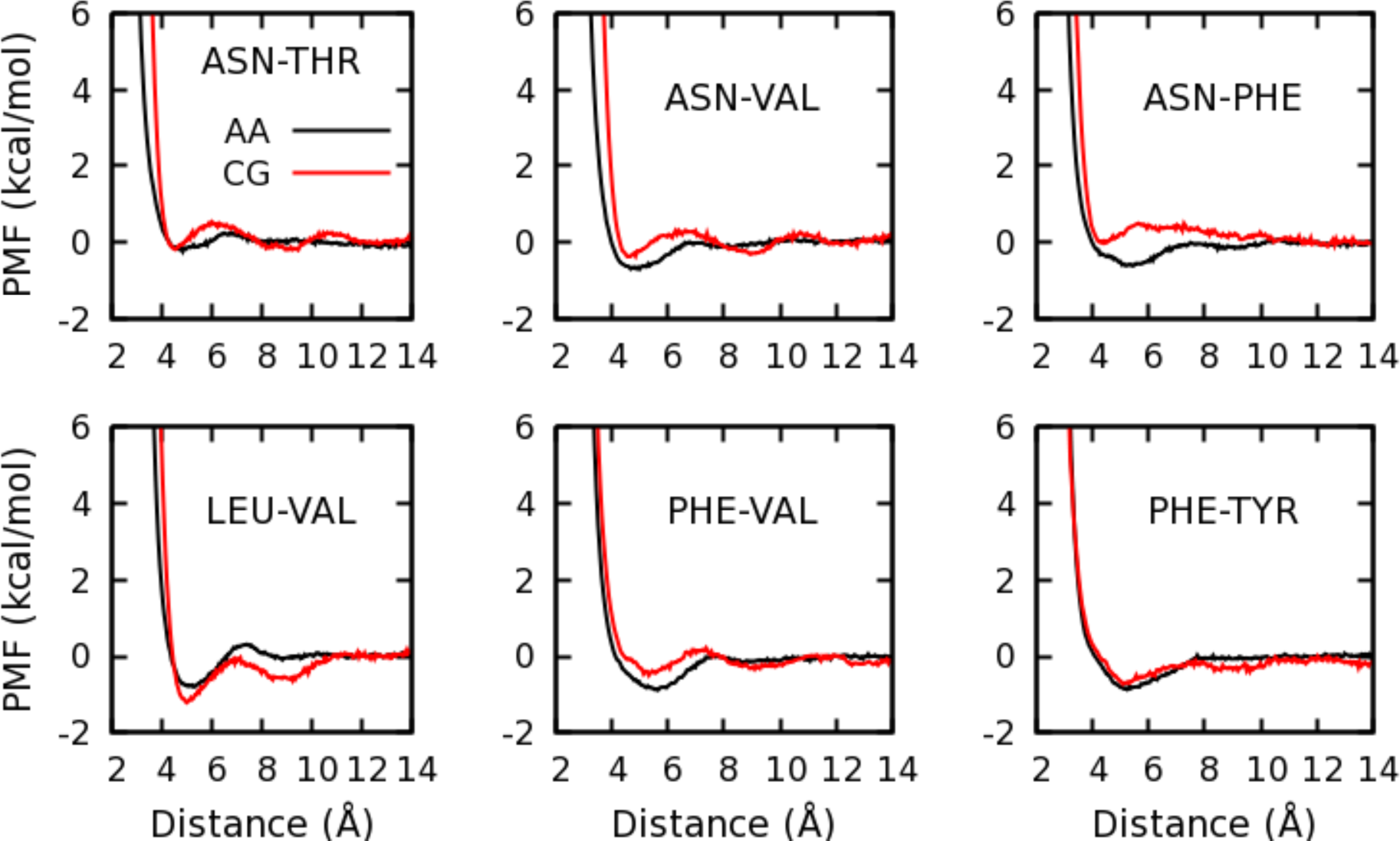
Dimerization free energies of side chain analogs. Black lines were calculated using CHARMM all-atom FF, and red lines were calculated using the initial LJ parameters employing the combination rules that are kept in the SPICA FF.

To optimize the LJ parameters between the side chains with multiple particles and to simultaneously optimize the interactions with water (see section 3.1), the RDF of side chain dimers in water from AA MD was used as a reference for charged and HIS side chains (target side chains). In the calculations, we put 12 target side-chain dimers and another 12 side-chain dimers were placed in water, to which Na^+^ or Cl^-^ ions were added for neutralization. Except for PRO, which shares the same side chain analog as VAL, there are a total of 17 kinds of side chains. Therefore, 85 systems were built, with each run lasting 10 ns. The last 5 ns (5,000 structures) were used for the analysis. The RDFs between the target side chain particles and all protein particles are shown in Figure S4. For most pairs, the AA RDFs (black lines) showed a single peak between 4-6 Å. The initial parameters employing combination rules largely underestimated the interactions for the target side chains, with the blue lines showing little or no peak between 4-6 Å. The present SPICA model greatly improved the RDF, although the first peak was slightly higher for some pairs. Notably, the RDFs converged to 1 at a distance of 15 Å for most pairs, suggesting that the choice of LJ cutoff was reasonable.

### 3.3 Free energy profile of SC analogs across DOPC bilayer

The free energy profiles of the SC analogs along the normal of the DOPC bilayer were used to optimize the CG parameters between SCs and lipid CG segments. The CG system consisted of 60 DOPC and 1,000 water, and a single SC analog was neutralized with Na^+^ or Cl^-^ when needed. In total, 25 umbrella windows were used in the range of 40 Å for umbrella sampling, and for each window, constraint MD runs were performed for 10-100 ns to obtain a well converged free energy profile using WHAM.^60^ The AA-MD results with the OPLS FF were used as reference data for the CG parameter optimization.^62^

Figure 3 shows the transfer free energy profiles of the 17 side-chain analogs from the DOPC lipid bilayer to water. The agreement between the AA and CG PMFs is excellent for polar and hydrophobic side chains, although the CG interface energy is slightly higher than that of AA for ASN and THR and slightly lower for CYS (black and red lines in the middle three layers of Figure 3). The CG results for the aromatic side chains are great (the bottom layer of Figure 3). The position of the energy minimum in the interface region shifted towards water for TYR and TRP in the CG MD. The difference in partitioning free energy between the AA and CG models was within 1 kcal/mol for most side chains, except for charged ones (the top layer of Figure 3). The free energy penalties for moving the charged side chains inside the lipid bilayer were underestimated by the CG model but were still very high (over 10 kcal/mol for ARG, ASP and GLU, and 4.6 kcal/mol for LYS). Therefore, the probability of entering the membrane was negligible for the charged residues. Overall, the SPICA FF can reproduce satisfactory partitioning between water and the lipid bilayer for amino acid side chain analogs.

**Figure 3.**
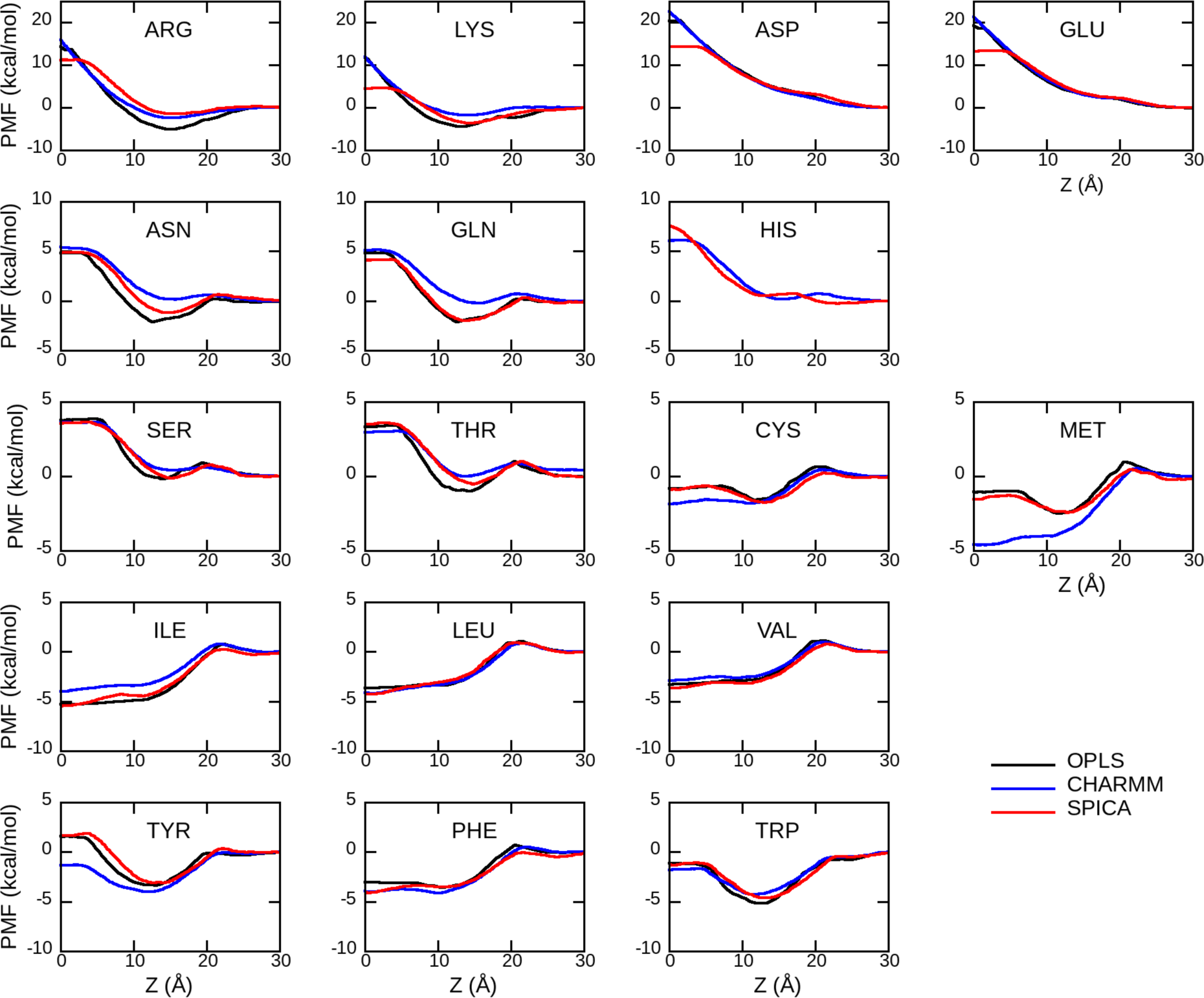
The transfer free energy profiles of side chain analogs along the DOPC bilayer normal. The distance between the centers of mass of the side chain analog and lipid bilayer along the normal of bilayer are shown along the x-axis. 0-10 Å corresponds to lipid tail region and 25-30 Å is the bulk water region.

For verification purposes, the AA PMFs were also calculated using the CHARMM36 FF and TIP3P water model (blue lines in Figure 3). This AA system included 72 DOPC and about 3750 water molecules. In contrast to other AA simulations in this study, the simulations were run using NAMD 2.11, and the PMFs were estimated by using adaptive biasing force (ABF).^63^ Langevin thermostat and Langevin-piston barostat^64^ were applied to control the temperature at 298 K and pressure of 1 atm, respectively. The same reaction coordinate was used as that in the CG MD and was split into six windows spanning from 0 to 30 Å with a spacing of 5 Å. Briefly, 200 ns MD simulation of the sampling was performed for each window. The total simulation time to estimate the free energy profile was 1.2 μs for each system. HIS was missing in the previous AA-MD, which confirms the reasonable agreement of the free energy profile of the HIS between SPICA and CHARMM FFs. As shown in Figure 3, the interfacial energy was slightly higher for the CHARMM FF than for the OPLS FF for the ARG, LYS, ASN, GLN, THR, and ILE side chains. According to the transfer free energy, SER and ILE were more hydrophilic and CYS, MET, and TYR were more hydrophobic in CHARMM FF than in OPLS FF. Regardless, we confirmed that the SPICA FF provides similar free energy profiles to those from the OPLS-FF; the deviation of each profile was similar to or less than the difference between the OPLS and the CHARMM-FF.

### 3.4 Penetration depths and tilt angles of alpha-helical peptides/proteins

We further optimized the LJ parameters between the backbone (BB) of peptides/proteins and water/lipid CG segments to reproduce the penetration depths and tilt angles of peripheral peptides/proteins obtained from the OPM database.^65^ The test set was taken from alpha-helical peptides/proteins in the OPM database, excluding ones with non-standard amino acid, acetylated N-terminus, amidated C-terminus, or disulfide bonds. In total, 97 peripheral alpha-helical peptides and 11 peripheral alpha-helical proteins were simulated. Initially, the peptides/proteins were placed approximately 2 Å away from the bilayer surface (128 DOPC molecules), solvated with water (3400-4800 molecules), and neutralized with Na^+^ or Cl^-^ ions (if necessary). After 1,000-steps energy minimization, the simulation run for 100−200 ns and the last 75 ns were used for analysis. To facilitate contact between peptides/proteins and lipids in a short time, a flat-bottomed potential was added to the centers of mass of peptides/proteins and lipids to limit their distance less than 10 Å far from the initial value.

Figure 4 shows the penetration depths and tilt angles of the 97 simulated peripheral peptides, together with those from the OPM database. The results for the 11 peripheral proteins are shown in Figure S5. Overall, the CG results agreed well with the OPM predictions. For the penetration depths, peptides 1fry, 1g92, 2k98, and 2kns exhibited rather large discrepancies between the CG and OPM results. In CG MD, cathelin-related peptide 1fry flipped up-to-down but maintained the correct interfacial interaction. Experiments show that both ends (top and bottom) have affinities to lipopolysaccharide.^66^ Poneratoxin 1g92 is a peptide that is deeply inserted into the lipid bilayer,^67^ although this insertion was only negligibly observed in CG MD. The agreement for tilt angles was better than for penetration depths, and only 5xes showed a large deviation from the OPM result. As a part of the TK9 peptide of the spike glycoprotein, 5xes is a short peptide with two alpha-helical rings, and the charged terminal backbone particles may be responsible for the flat orientation in CG MD. In Figure 4, we selected an alpha-helical peptide to parameterize the BB interaction parameters, although we also examined the same parameters for beta-harpin peptides. The data are presented in Figure S6 for comparison with the OPM database. Even though these peptides were not included in the parameter fitting, the CG-MD results were in reasonable agreement with the OPM database in terms of the anchoring depth and orientation of most peptides.

**Figure 4.**
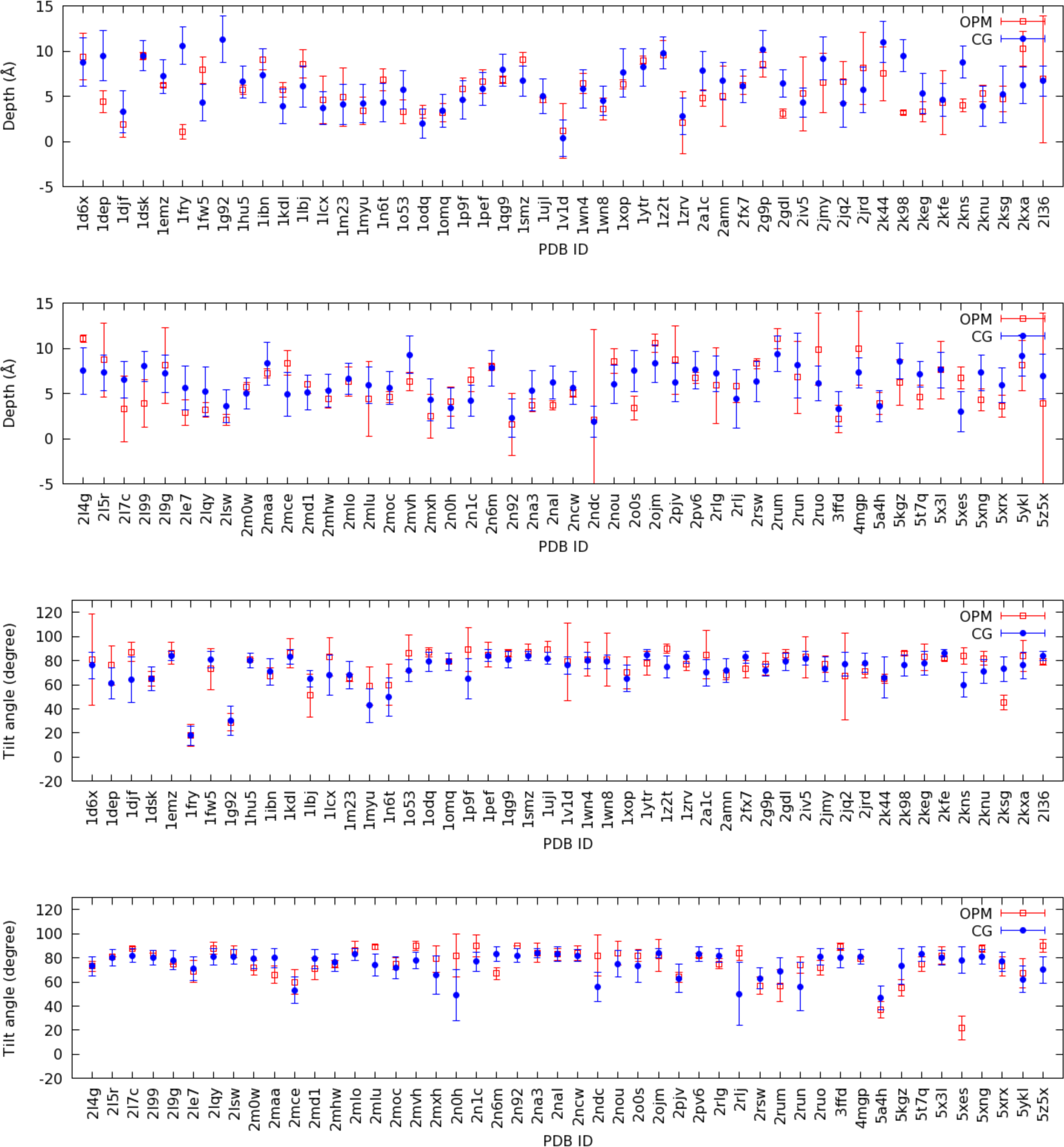
Penetration depths and tilt angles of peripheral peptides to DOPC membranes.

The tilt angle for the transmembrane peptide WALP was also calculated to evaluate the interaction between the backbone and water/lipid. WALP (sequence: GWW(LA)_8_LWWA) was inserted into the membrane (128 lipid molecules) at the beginning and then solvated with 2210 water molecules. The simulations were run for 1 µs, and the last 750 ns was used for analysis.

As shown in Table 4, the CG results are consistent with the experimental and previous MD simulation results, considering the large deviations of the CG calculations. The tilt angle in the DMPC bilayer was larger than that in the DOPC bilayer because of the hydrophobic matching in the thinner DMPC membrane. The tilt angle was slightly overestimated in the SPICA FF, although it was not easy to precisely compare the tilt angles due to the large thermal fluctuations.

**Table 4.**
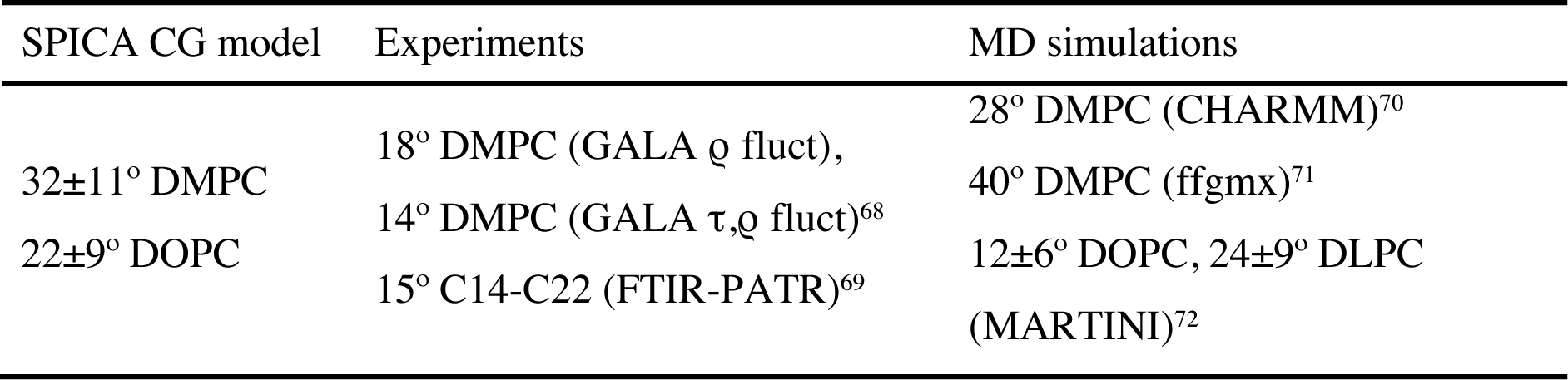
Tilt angle of transmembrane peptide WALP

### 3.5 Dimerization free energy of transmembrane helices

The dimerization free energies of transmembrane helices were used to optimize the LJ parameters for the BB−BB and BB−SC segments. For this test, we chose five different transmembrane helices, that is, GpA, SerZip, WALP peptides, and the transmembrane segments of EphA1 and ErbB1. The dimerization free energies were calculated along the distance between the centers of mass of two transmembrane helices by using umbrella sampling and WHAM methods.^60^ For GpA, two parallel helices were placed into a system containing 123 DPPC and 2553 water. The two SerZip helices were also parallel and surrounded by 128 POPC and 2046 CG water. Two antiparallel WALP helices were inserted into the lipid bilayer of 128 DOPC and solvated with 2210 CG water. The transmembrane segments of EphA1 and ErbB1 were embedded in 254 DMPC and 4197 CG water, and in 255 DLPC and 4050 CG water, respectively. In all cases, counter CG ions (Na^+^ or Cl^-^) were added to neutralize the total system. Peptides were stabile as dimers in these lipid membranes in an equilibrium CG MD simulation. We pulled the two helices apart to generate the initial configurations for the dimerization free energy calculation using colvars.^73^ To accelerate the convergence, ABF simulations were performed for separating windows with a width of ∼4 Å. Then, 5 µs-long MD was run for each window of each system for the analysis.

Table 5 shows the calculated dimerization free energies using the present SPICA FF and other FF, as well as the experimental data. For GpA, the calculated dimerization free energy in DPPC was calculated as -3.9 ± 0.5 kcal/mol using the present SPICA FF, which was within the range of experimental values (Table 5). The dimerization free energy of SerZip estimated with the SPICA FF agrees well with the high binding affinity identified in POPC by sedimentation equilibrium experiments using ultracentrifugation (Table 5). For WALP, however, the dimerization free energy of the SPICA FF seemed to be overestimated, compared to the experimental estimation by Yano and Matsuzaki.^69^ The sequence used in the experiments was (AALALAA)_3_, and the repetition of ALA and LEU in the hydrophobic region was similar to the sequence of WALP (GWW(LA)_8_LWWA) used in this work, although it could be a reason for the deviation. The dimerization free energy for EphA1 in the present CG FF showed excellent agreement with the experimental value. The same is true for ErbB1 dimerization free energy. In summary, although the agreement between the CG MD results and experimental data was conditional, the dimerization free energies of the five peptides could be reasonably reproduced by scaling the LJ parameters ɛ of the BB-BB and BB-SC segments.

**Table 5.** Dimerization free energies of transmembrane helices

|  | SPICA CG model<br>(kcal/mol) | Experiments (kcal/mol) | MD simulations (kcal/mol) |
| --- | --- | --- | --- |
| GpA | -3.9 ± 0.5 DPPC | -3.7~-7.5 C<sub>10</sub>~C<sub>12</sub><br>detergent micelles<sup>74</sup><br>-7.0 C<sub>8</sub>E<sub>5</sub> micelles<sup>75</sup><br>-3.2~-4.0 cell<br>membranes<sup>76,77</sup> | -11.5 ± 0.4 dodecane slab<br>(CHARMM27)<sup>78</sup><br>-9 DPPC (MARTINI2.1)<sup>79</sup><br>~ -6 (MARTINI3)<sup>28</sup> |
| SerZip | -7.4 ± 0.7 POPC | -4.5 DLPC, <-8 POPC<sup>80</sup> | |
| WALP | -10.1 ± 0.9 DOPC | -2.2±0.2~-6.2±0.3<br>C14~C22<br>dimonounsaturated<br>phosphocholine lipid<br>bilayers<sup>69</sup> | -8 DMPC, -6 DPPC, -5 DOPC<br>(antiparallel, MARTINI)<sup>81</sup><br>~ -3.1 (MARTINI3)<sup>28</sup> |
| EphA1 | -3.6 ± 0.6 DMPC | -3.7 ± 0.1 DMPC<sup>82</sup> | -14.3 ± 0.1 DPPC (MARTINI)<sup>83</sup><br>-7.1 ± 0.2 ~ -8.0 ± 0.2 DMPC<br>(MARTINI)<sup>84</sup><br>~ -5.2 (MARTINI3)<sup>28</sup> |
| ErbB1 | -2.5 ± 0.4 DLPC | -2.5 ± 0.1 DLPC<sup>85</sup> | -9 ± 1 DPPC (MARTINI)<sup>86</sup><br>-9.4 ± 0.2 DLPC (MARTINI)<sup>84</sup><br>~ -2.6 (MARTINI3)<sup>28</sup> |

## 4. APPLICATION STUDIES

### 4.1 Prediction of native structures from decoy set

To evaluate the accuracy of the SPICA FF, we first evaluated whether the native structures could be selected from the decoy sets that contain structures generated using various techniques. As in our previous work,^40^ five decoy sets (*4state_reduced*, *fisa*, *fisa_casp3*, *lmds*, and *lattice_ssfit*), provided by Decoys R Us, were used.^87–91^ The model succeeded in predicting the native structures when the rank was 1. As shown in Table 6, the SPICA FF showed a better performance than the previous model (SDK2009)^40^ in decoy set *lmds*, similar in decoy set *fisa* and *fisa_casp3*, and worse in decoy set *4state_reduced* and *lattice_ssfit*. Both the SDK2009 and SPICA FF yield the better scores than the MARTINI FF (2.2) in predicting the native structures. (We have not succeeded to use MARTINI3 FF for this test).

**Table 6.** Rank of native structures in decoy sets

| Decoy set | PDB ID | SDK2009<sup>40</sup> | SPICA | MARTINI2.2<sup>27</sup> |
| --- | --- | --- | --- | --- |
| *4state_reduced* | 1ctf | 1 | 83 | 144 |
| *4state_reduced* | 1r69 | 1 | 12 | 73 |
| *4state_reduced* | 1sn3 | 151 | 99 | 657 |
| *4state_reduced* | 2cro | 1 | 30 | 48 |
| *4state_reduced* | 3icb | 387 | 226 | 148 |
| *4state_reduced* | 4pti | 267 | 90 | 672 |
| *4state_reduced* | 4rxn | 59 | 3 | 133 |
| *fisa* | 1fc2 | 64 | 12 | 382 |
| *fisa* | 1hdd-C | 3 | 8 | 469 |
| *fisa* | 2cro | 1 | 1 | 333 |
| *fisa* | 4icb | 1 | 1 | 453 |
| *fisa_casp3* | 1bg8-A | 1 | 1 | 315 |
| *fisa_casp3* | 1bl0 | 1 | 1 | 1 |
| *fisa_casp3* | 1eh2 | 6 | 42 | 356 |
| *fisa_casp3* | 1jwe | 1 | 1 | 1 |
| *fisa_casp3* | smd3 | 1 | 1 | 2 |
| *lmds* | 1b0n-B | 303 | 67 | 32 |
| *lmds* | 1ctf | 1 | 1 | 227 |
| *lmds* | 1dtk | 155 | 7 | 200 |
| *lmds* | 1fc2 | 1 | 1 | 1 |
| *lmds* | 1igd | 3 | 3 | 1 |
| *lmds* | 1shf-A | 2 | 1 | 234 |
| *lmds* | 2cro | 1 | 1 | 39 |
| *lmds* | 2ovo | 258 | 1 | 1 |
| *lattice_ssfit* | 1beo | 19 | 24 | 1661 |
| *lattice_ssfit* | 1ctf | 1 | 8 | 277 |
| *lattice_ssfit* | 1dkt-A | 1 | 1 | 685 |
| *lattice_ssfit* | 1fca | 1 | 4 | 1700 |
| *lattice_ssfit* | 1nkl | 1 | 167 | 1128 |
| *lattice_ssfit* | 1pgb | 1 | 777 | 1886 |
| *lattice_ssfit* | 1trl-A | 1 | 6 | 198 |
| *lattice_ssfit* | 4icb | 1 | 1 | 15 |

### 4.2 Assembly of membrane proteins in DOPC bilayer

The assembly of membrane proteins in a lipid membranes has been extensively investigated because of their importance in cell metabolism and gene transcription. Here, we examined the stability of the protein assembly in the DOPC membrane to evaluate the SPICA FF. Twenty-six membrane protein assemblies consisting of three or four subunits were selected from the OPM database for these test calculations.^46^ For ease of simulation, all extracellular and juxtamembrane domains were omitted. MD simulations were carried out for 500 ns, and the average RMSD of the backbone (as a membrane protein assembly) during the last 100 ns was calculated and compared to the initial structures. As shown in Figure 5, the overall structures were well stabilized throughout the simulations with small RMSDs of less than 5 Å, except in a few cases. The large RMSD for voltage-dependent potassium channel KvAP (2a0l)^92^ was caused by the movement of peripheral voltage-sensing domains, while the large RMSD for mechanosensitive channels of large (MscL) conductances (3hzq)^93^ resulted from the high flexibility of loops.

**Figure 5.**
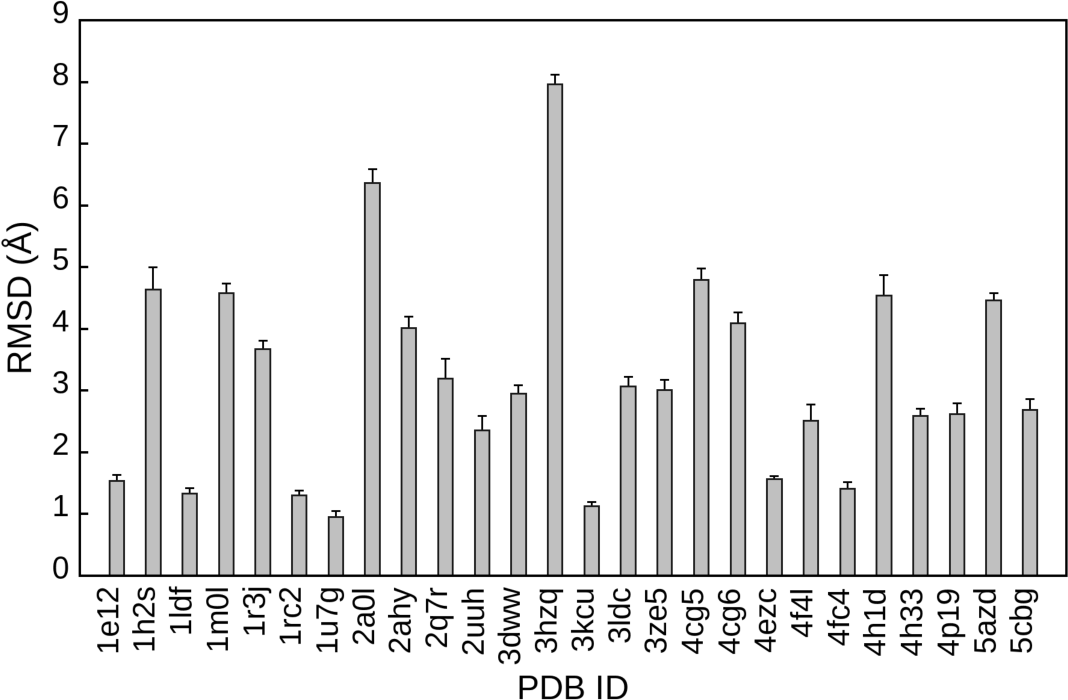
RMSDs of membrane protein assembly in DOPC bilayer. The average RMSD of backbone during the last 100 ns was calculated with respect to the initial structures.

### 4.3 Lipid translocation through scramblase

The phospholipid flip-flop is an important process for maintaining the asymmetric lipid composition of the cellular membrane, which plays a crucial role in biological functions^94^. Because the transverse movement of phospholipids is hampered by the hydrophobic core of the lipid bilayer, it is catalyzed by enzymes, such as flippase, floppase, and scramblase. The translocation mechanism of scramblase is of particular interest. Recent studies have shown that phospholipid translocation occurs at the hydrophilic cavities on scramblase surfaces.^95, 96^ As the pathway of lipid translocation requires a correct description of the hydrophobicity of amino acids, we assessed our protein model by simulating lipid translocation with scramblase. In our previous CG MD simulations using the SPICA FF, no phospholipid flip-flop was observed in the PC bilayer membrane in the absence of scramblase. We embedded nhTMEM16 (PDB ID: 4WIS) in the POPC membrane and ran 3-μs CG MD simulations. Figure 6A shows a snapshot of the selected segments of the simulated system with the protein surface and phosphates as spheres, demonstrating the membrane deformation due to the insertion of scramblase and the phosphate groups of flipping lipids in the hydrophilic cavities on the protein surface. Except for the hydrophilic cavities, we did not observe any transverse motion of the lipid. We also measured the number of lipid molecules during the 3-μs MD simulation. The phospholipids in the cavities within 20-Å thickness were occupied by 3.48 ± 0.747 and 4.11 ± 0.857 lipids on average (Figure 6B), which is consistent with the experimental results.^97^ Only one full translocation occurred during the simulation. The simulation result of scramblase shows that the hydrophobicity of the present SPICA FF was correctly modeled.

**Figure 6.**
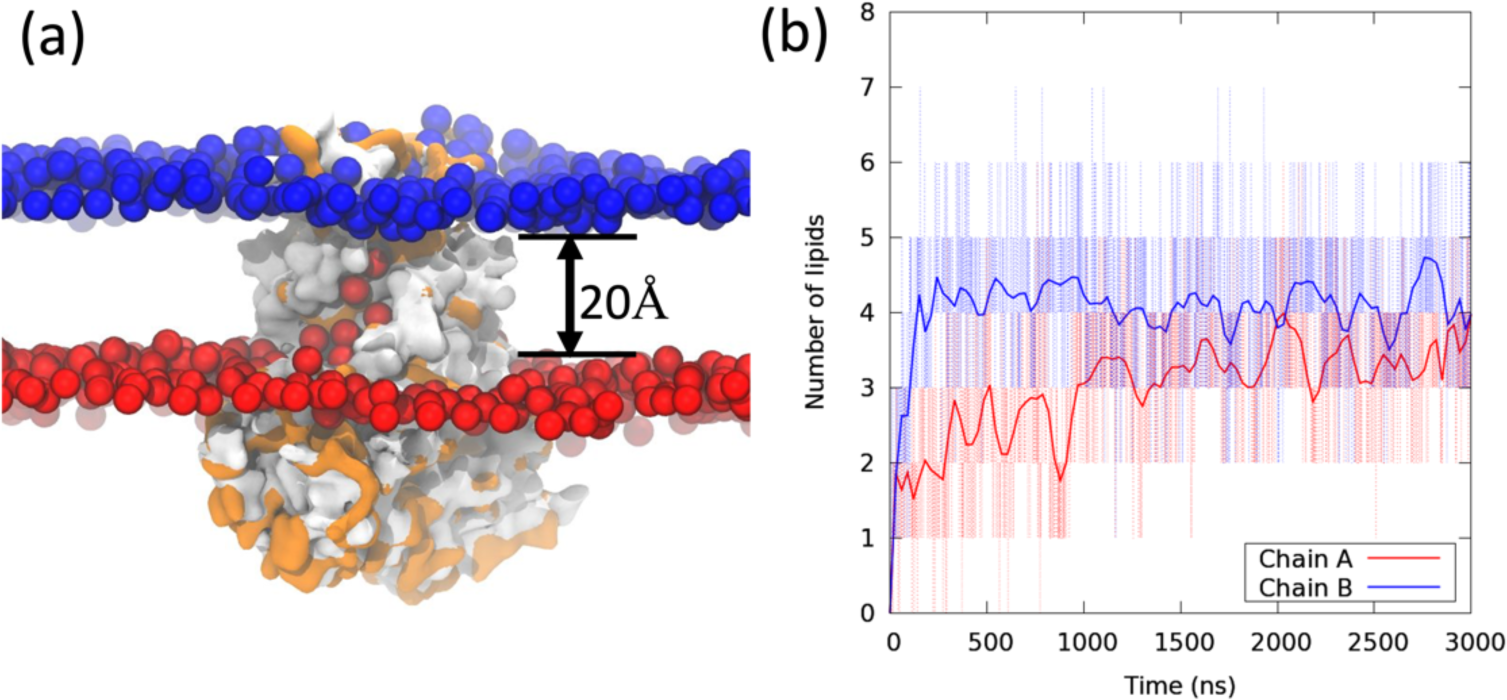
Results of the CG MD simulation of TMEM16. (a) Snapshot of hydrophilic grooves of scramblase. Orange surfaces show hydrophilic amino acids. Beads and surface represent phosphate group of lipid (bead type: PH) and scramblase, respectively. Red and blue beads represent the PH presented in the lower and upper leaflets, respectively. (b) Number of lipids in the hydrophilic cavity within a thickness of 20 Å. Red and blue lines denote the number of lipids from each leaflet in each chain.

### 4.4 Stability of poliovirus capsid in water

Although the current SPICA FF was mainly designed for simulations of membrane proteins, we also examined its performance for a protein assembly. Poliovirus is the causative agent of poliomyelitis. Its icosahedral capsid contains four kinds of proteins (VP1, VP2, VP3, and VP4), each of which has 60 copies. AA MD of poliovirus has been performed in a previous work.^98^ For ease of comparison, we took the equilibrated atomic structure as initial structure and then converted it to CG system (with 619157 water, 5446 Na^+^ and 5206 Cl^-^). ENM was applied only to the intra-chain backbone particles. A CG-MD simulation was carried out for 1 µs to confirm the stability of the capsid (Figure 7a). The average radius of poliovirus capsid during the last 500 ns was about 132.8 Å in CG MD, which agreed well with 133.6 Å in AA MD. RMSF was also compared between the AA and CG MD. As shown in Figure 7b, the CG model was slightly more flexible than the AA model, which may be improved by changing the ENM parameters. Because there was no ENM between the inter-chain CG particles, the interaction between chains was dominated by non-bonded interactions. The stable capsid suggests that the SPICA FF can also be used for simulations of soluble proteins in water.

**Figure 7.**
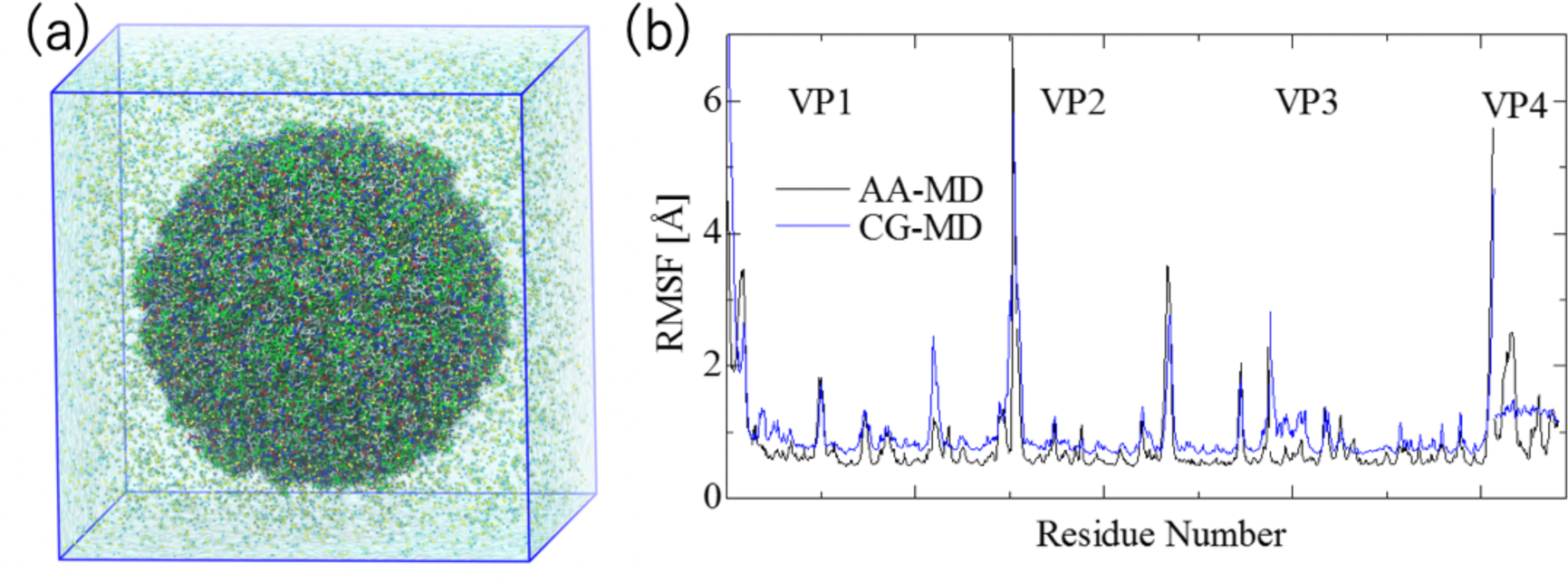
(a) A CG snapshot of poliovirus in water. (b) Comparison of RMSFs of poliovirus from AA MD and CG MD.

## 5. CONCLUSIONS

In this study, we developed a CG protein model compatible with the SPICA lipid and water model. This CG protein model was designed for membrane proteins to investigate behaviors, such as the assembly of proteins in the lipid bilayer and the deformation of the lipid bilayer by membrane proteins. Bonded parameters were created from structures of the PDB database or AA MD, and secondary structures were maintained using the ENM method. Non-bonded parameters were optimized by reproducing multiple properties from both the experiments and the AA MDs. The resulting CG protein model could reproduce: (i) a stable structure of membrane protein assembly in the lipid bilayer, (ii) a reasonable interaction between protein and lipid/water (partitioning of SC in lipid/water, depth and tilt angle of membrane peptide/protein), and (iii) a reasonable interaction between protein and protein (dimerization free energy of SCs in water, dimerization free energy of transmembrane helices). The present SPICA model has been already used in a recent study of the protein seipin, which induces the lipid droplet formation.^99^ Although this CG model was designed for membrane proteins, it can also reproduce the stable structure of the poliovirus capsid in aqueous solution. The application of ENM is limited to studies involving large conformational changes, such as protein folding. In other applications, such as the prediction of native structures and lipid translocation, the use of the CG model is more suitable. Overall, the present SPICA CG protein model is a suitable choice for the simulation of membrane proteins.

Detailed information of the SPICA force field, including all the parameters and simulation tools used to prepare the initial setups for MD simulations, is freely available from the webpage (https://www.spica-ff.org/).

## ACKNOWLEDGMENTS

The authors thank Prof. S. Okazaki and Dr. Y. Andoh for sharing their all-atom configuration of polio virus capsid. This work was supported by JSPS KAKENHI (grant no. 21H01880), and MEXT as “Program for Promoting Researches on the Supercomputer Fugaku” (Biomolecular dynamics and function in a living cell using atomistic and coarse-grained simulations) (no. JPMXP1020200101), and by the US National Science Foundation grant CHE-1212416. Calculations were performed at the facilities of the Research Center for Computational Science, Okazaki, the Institute for Solid State Physics, and the University of Tokyo and, in part, on the Fugaku computer hosted at the RIKEN Advanced Institute for Computational Science (proposal no. hp200135 and hp210177).

## REFERENCES

(1) Shinoda, W.; Devane, R.; Klein, M. L. Multi-Property Fitting and Parameterization of a Coarse Grained Model for Aqueous Surfactants. Mol. Simul. 2007, 33 (1–2), 27–36.

(2) Basdevant, N.; Borgis, D.; Ha-Duong, T. Modeling Protein-Protein Recognition in Solution Using the Coarse-Grained Force Field SCORPION. J. Chem. Theory Comput. 2013, 9 (1), 803–813.

(3) Li, M.; Zhang, J. Z. H. Two-Bead Polarizable Water Models Combined with a Two-Bead Multipole Force Field (TMFF) for Coarse-Grained Simulation of Proteins. Phys. Chem. Chem. Phys. 2017, 19 (10), 7410–7419.

(4) Frezza, E.; Lavery, R. Internal Normal Mode Analysis (INMA) Applied to Protein Conformational Flexibility. J. Chem. Theory Comput. 2015, 11 (11), 5503–5512.

(5) Cheon, M.; Chang, I.; Hall, C. K. Extending the PRIME Model for Protein Aggregation to All 20 Amino Acids. Proteins Struct. Funct. Bioinforma. 2010, 78 (14), 2950–2960.

(6) Emperador, A.; Sfriso, P.; Villarreal, M. A.; Gelpí, J. L.; Orozco, M. PACSAB: Coarse-Grained Force Field for the Study of Protein-Protein Interactions and Conformational Sampling in Multiprotein Systems. J. Chem. Theory Comput. 2015, 11 (12), 5929–5938.

(7) Frembgen-Kesner, T.; Andrews, C. T.; Li, S.; Ngo, N. A.; Shubert, S. A.; Jain, A.; Olayiwola, O. J.; Weishaar, M. R.; Elcock, A. H. Parametrization of Backbone Flexibility in a Coarse-Grained Force Field for Proteins (COFFDROP) Derived from All-Atom Explicit-Solvent Molecular Dynamics Simulations of All Possible Two-Residue Reptides. J. Chem. Theory Comput. 2015, 11 (5), 2341–2354.

(8) da Silva, F. L.; Sterpone, F.; Derreumaux, P. OPEP6: A New Constant-PH Molecular Dynamics Simulation Scheme with OPEP Coarse-Grained Force Field. J. Chem. Theory Comput. 2019, 15 (6), 3875–3888.

(9) Li, W.; Wang, W.; Takada, S. Energy Landscape Views for Interplays among Folding, Binding, and Allostery of Calmodulin Domains. Proc. Natl. Acad. Sci. U. S. A. 2014, 111 (29), 10550–10555.

(10) Sanyal, T.; Mittal, J.; Shell, M. S. A Hybrid, Bottom-up, Structurally Accurate, Go-like Coarse-Grained Protein Model. J. Chem. Phys. 2019, 151 (4), 044111.

(11) Kapoor, A.; Travesset, A. Folding 19 Proteins to Their Native State and Stability of Large Proteins from a Coarse-Grained Model. Proteins Struct. Funct. Bioinforma. 2014, 82 (3), 505–516.

(12) Liu, X.; Chen, J. HyRes: A Coarse-Grained Model for Multi-Scale Enhanced Sampling of Disordered Protein Conformations. Phys. Chem. Chem. Phys. 2017, 19 (48), 32421–32432.

(13) Dawid, A. E.; Gront, D.; Kolinski, A. SURPASS Low-Resolution Coarse-Grained Protein Modeling. J. Chem. Theory Comput. 2017, 13 (11), 5766–5779.

(14) Davtyan, A.; Simunovic, M.; Voth, G. A. Multiscale Simulations of Protein-Facilitated Membrane Remodeling. J. Struct. Biol. 2016, 196 (1), 57–63.

(15) Bereau, T.; Wang, Z.-J.; Deserno, M. More than the Sum of Its Parts: Coarse-Grained Peptide-Lipid Interactions from a Simple Cross-Parametrization. J. Chem. Phys. 2014, 140 (11), 115101.

(16) Hills, R. D. Refining Amino Acid Hydrophobicity for Dynamics Simulation of Membrane Proteins. PeerJ 2018, 6, e4230.

(17) Tesei, G.; Vazdar, M.; Lund, M. Coarse-Grained Model of Titrating Peptides Interacting with Lipid Bilayers. J. Chem. Phys. 2018, 149 (24), 244108.

(18) Shen, H.; Li, Y.; Xu, P.; Li, X.; Chu, H.; Zhang, D.; Li, G. An Anisotropic Coarse-Grained Model Based on Gay-Berne and Electric Multipole Potentials and Its Application to Simulate a DMPC Bilayer in an Implicit Solvent Model. J. Comput. Chem. 2015, 36 (15), 1103–1113.

(19) Kar, P.; Gopal, S. M.; Cheng, Y. M.; Panahi, A.; Feig, M. Transferring the PRIMO Coarse-Grained Force Field to the Membrane Environment: Simulations of Membrane Proteins and Helix-Helix Association. J. Chem. Theory Comput. 2014, 10 (8), 3459– 3472.

(20) Vorobyov, I.; Kim, I.; Chu, Z. T.; Warshel, A. Refining the Treatment of Membrane Proteins by Coarse-Grained Models. Proteins Struct. Funct. Bioinforma. 2015, 84 (1), 92–117.

(21) Kim, B. L.; Schafer, N. P.; Wolynes, P. G. Predictive Energy Landscapes for Folding α-Helical Transmembrane Proteins. Proc. Natl. Acad. Sci. U. S. A. 2014, 111 (30), 11031–11036.

(22) Pulawski, W.; Jamroz, M.; Kolinski, M.; Kolinski, A.; Kmiecik, S. Coarse-Grained Simulations of Membrane Insertion and Folding of Small Helical Proteins Using the CABS Model. J. Chem. Inf. Model. 2016, 56 (11), 2207–2215.

(23) Ziȩba, K.; Ślusarz, M.; Ślusarz, R.; Liwo, A.; Czaplewski, C.; Sieradzan, A. K. Extension of the UNRES Coarse-Grained Force Field to Membrane Proteins in the Lipid Bilayer. J. Phys. Chem. B 2019, 123 (37), 7829–7839.

(24) Alford, R. F.; Koehler Leman, J.; Weitzner, B. D.; Duran, A. M.; Tilley, D. C.; Elazar, A.; Gray, J. J. An Integrated Framework Advancing Membrane Protein Modeling and Design. PLoS Comput. Biol. 2015, 11 (9), e1004398.

(25) Kim, Y. C.; Hummer, G. Coarse-Grained Models for Simulations of Multiprotein Complexes: Application to Ubiquitin Binding. J. Mol. Biol. 2008, 375 (5), 1416–1433.

(26) Marrink, S. J.; Risselada, H. J.; Yefimov, S.; Tieleman, D. P.; De Vries, A. H. The MARTINI Force Field: Coarse Grained Model for Biomolecular Simulations. J. Phys. Chem. B 2007, 111 (27), 7812–7824.

(27) De Jong, D. H.; Singh, G.; Bennett, W. F. D.; Arnarez, C.; Wassenaar, T. A.; Schäfer, L. V; Periole, X.; Tieleman, D. P.; Marrink, S. J. Improved Parameters for the Martini Coarse-Grained Protein Force Field. J. Chem. Theory Comput. 2013, 9 (1), 687–697.

(28) Souza, P. C. T.; Alessandri, R.; Barnoud, J.; Thallmair, S.; Faustino, I.; Grünewald, F.; Patmanidis, I.; Abdizadeh, H.; Bruininks, B. M. H.; Wassenaar, T. A.; Kroon, P. C.; Melcr, J.; Nieto, V.; Corradi, V.; Khan, H. M.; Domański, J.; Javanainen, M.; Martinez-Seara, H.; Reuter, N.; Best, R. B.; Vattulainen, I.; Monticelli, L.; Periole, X.; Tieleman, D. P.; Vries, A. H. de; Marrink, S. J. Martini 3: A General Purpose Force Field for Coarse-Grained Molecular Dynamics. Nat. Methods 2021, 18 (4), 382–388.

(29) Han, W.; Wan, C.-K.; Jiang, F.; Wu, Y.-D. PACE Force Field for Protein Simulations. Full Parameterization of Version 1 and Verification. J. Chem. Theory Comput. 2010, 6 (11), 3373–3389.

(30) Han, W.; Wan, C.-K.; Wu, Y.-D. PACE Force Field for Protein Simulations. 2. Folding Simulations of Peptides. J. Chem. Theory Comput. 2010, 6 (11), 3390–3402.

(31) Wan, C. K.; Han, W.; Wu, Y. D. Parameterization of PACE Force Field for Membrane Environment and Simulation of Helical Peptides and Helix-Helix Association. J. Chem. Theory Comput. 2012, 8 (1), 300–313.

(32) Han, W.; Schulten, K. Further Optimization of a Hybrid United-Atom and Coarse-Grained Force Field for Folding Simulations: Improved Backbone Hydration and Interactions between Charged Side Chains. J. Chem. Theory Comput. 2012, 8 (11), 4413–4424.

(33) Darré, L.; Machado, M. R.; Brandner, A. F.; González, H. C.; Ferreira, S.; Pantano, S. SIRAH: A Structurally Unbiased Coarse-Grained Force Field for Proteins with Aqueous Solvation and Long-Range Electrostatics. J. Chem. Theory Comput. 2015, 11 (2), 723–739.

(34) Barrera, E. E.; Machado, M. R.; Pantano, S. Fat SIRAH: Coarse-Grained Phospholipids to Explore Membrane-Protein Dynamics. J. Chem. Theory Comput. 2019, 15 (10), 5674–5688.

(35) Shinoda, W.; Devane, R.; Klein, M. L. Zwitterionic Lipid Assemblies: Molecular Dynamics Studies of Monolayers, Bilayers, and Vesicles Using a New Coarse Grain Force Field. J. Phys. Chem. B 2010, 114 (20), 6836–6849.

(36) Seo, S.; Shinoda, W. SPICA Force Field for Lipid Membranes: Domain Formation Induced by Cholesterol. J. Chem. Theory Comput. 2019, 15 (1), 762–774.

(37) Shinoda, W.; Nakamura, T.; Nielsen, S. O. Free Energy Analysis of Vesicle-to-Bicelle Transformation. Soft Matter 2011, 7 (19), 9012–9020.

(38) Kawamoto, S.; Nakamura, T.; Nielsen, S. O.; Shinoda, W. A Guiding Potential Method for Evaluating the Bending Rigidity of Tensionless Lipid Membranes from Molecular Simulation. J. Chem. Phys. 2013, 139 (3), 034108.

(39) Nakamura, T.; Shinoda, W. Method of Evaluating Curvature-Dependent Elastic Parameters for Small Unilamellar Vesicles Using Molecular Dynamics Trajectory. J. Chem. Phys. 2013, 138 (12), 124903.

(40) Devane, R.; Shinoda, W.; Moore, P. B.; Klein, M. L. Transferable Coarse Grain Nonbonded Interaction Model for Amino Acids. J. Chem. Theory Comput. 2009, 5 (8), 2115–2124.

(41) Tirion, M. M. Large Amplitude Elastic Motions in Proteins from a Single-Parameter, Atomic Analysis. Phys. Rev. Lett. 1996, 77 (9), 1905–1908.

(42) Shinoda, W.; Devane, R.; Klein, M. L. Coarse-Grained Force Field for Ionic Surfactants. Soft Matter 2011, 7 (13), 6178–6186.

(43) Frenkel, D.; Smit, B. Appendix D – Statistical Errors. *Underst*. Mol. Simul. 2002.

(44) Periole, X.; Cavalli, M.; Marrink, S.-J.; Ceruso, M. A. Combining an Elastic Network With a Coarse-Grained Molecular Force Field: Structure, Dynamics, and Intermolecular Recognition. J. Chem. Theory Comput. 2009, 5 (9), 2531–2543.

(45) Devane, R.; Klein, M. L.; Chiu, C.; Nielsen, S. O.; Shinoda, W.; Moore, P. B. Coarse-Grained Potential Models for Phenyl-Based Molecules: I. Parametrization Using Experimental Data. J. Phys. Chem. B 2010, 114 (19), 6386–6393.

(46) Lomize, M. A.; Pogozheva, I. D.; Joo, H.; Mosberg, H. I.; Lomize, A. L. OPM Database and PPM Web Server: Resources for Positioning of Proteins in Membranes. Nucleic Acids Res. 2012, 40 (Database issue), D370--D376.

(47) Plimpton, S. Fast Parallel Algorithms for Short-Range Molecular Dynamics. J. Comput. Phys. 1995, 117, 1–19.

(48) Parrinello, M.; Rahman, A. Polymorphic Transitions in Single Crystals: A New Molecular Dynamics Method. J. Appl. Phys. 1981, 52 (12), 7182–7190.

(49) Shinoda, W.; Shiga, M.; Mikami, M. Rapid Estimation of Elastic Constants by Molecular Dynamics Simulation under Constant Stress. Phys. Rev. B 2004, 69 (13), 134103.

(50) Nosé, S. A Unified Formulation of the Constant Temperature Molecular Dynamics Methods. J. Chem. Phys. 1984, 81, 511–519.

(51) Hoover, W. G. Canonical Dynamics: Equilibrium Phase-Space Distributions. Phys. Rev. A 1985, 31 (3), 1695–1697.

(52) Griffiths, M. Z.; Shinoda, W. TSPICA: Temperature- and Pressure-Dependent Coarse-Grained Force Field for Organic Molecules. J. Chem. Inf. Model. 2019, 59 (9), 3829– 3838.

(53) Abraham, M. J.; Murtola, T.; Schulz, R.; Páll, S.; Smith, J. C.; Hess, B.; Lindah, E. Gromacs: High Performance Molecular Simulations through Multi-Level Parallelism from Laptops to Supercomputers. SoftwareX 2015, 1-2, 19–25.

(54) Klauda, J. B.; Venable, R. M.; Freites, J. A.; O’Connor, J. W.; Tobias, D. J.; Mondragon-Ramirez, C.; Vorobyov, I.; MacKerell, A. D.; Pastor, R. W. Update of the CHARMM All-Atom Additive Force Field for Lipids: Validation on Six Lipid Types. J. Phys. Chem. B 2010, 114 (23), 7830–7843.

(55) Best, R. B.; Zhu, X.; Shim, J.; Lopes, P. E. M.; Mittal, J.; Feig, M.; MacKerell, A. D. Optimization of the Additive CHARMM All-Atom Protein Force Field Targeting Improved Sampling of the Backbone φ, ψ and Side-Chain Χ1 and Χ2 Dihedral Angles. J. Chem. Theory Comput. 2012, 8 (9), 3257–3273.

(56) Darden, T.; York, D.; Pedersen, L. Particle Mesh Ewald: An N·log(N) Method for Ewald Sums in Large Systems. J. Chem. Phys. 1993, 98 (12), 10089–10092.

(57) Essmann, U.; Perera, L.; Berkowitz, M. L.; Darden, T.; Lee, H.; Pedersen, L. G. A Smooth Particle Mesh Ewald Method. J. Chem. Phys. 1995, 103 (19), 8577–8593.

(58) Hess, B. P-LINCS: A Parallel Linear Constraint Solver for Molecular Simulation. J. Chem. Theory Comput. 2008, 4 (1), 116–122.

(59) Torrie, G. M.; Valleau, J. P. Nonphysical Sampling Distributions in Monte Carlo Free-Energy Estimation: Umbrella Sampling. J. Comput. Phys. 1977, 23 (2), 187–199.

(60) Kumar, S.; Rosenberg, J. M.; Bouzida, D.; Swendsen, R. H.; Kollman, P. A. THE Weighted Histogram Analysis Method for Free-energy Calculations on Biomolecules. I. The Method. J. Comput. Chem. 1992, 13 (8), 1011–1021.

(61) Wolfenden, R.; Andersson, L.; Cullis, P. M.; Southgate, C. C. B. Affinities of Amino Acid Side Chains for Solvent Water. Biochemistry 1981, 20 (4), 849–855.

(62) MacCallum, J. L.; Drew Bennett, W. F.; Peter Tieleman, D. Distribution of Amino Acids in a Lipid Bilayer from Computer Simulations. Biophys. J. 2008, 94 (9), 3393– 3404.

(63) Darve, E.; Pohorille, A. Calculating Free Energies Using Average Force Adaptive Biasing Force Method for Scalar and Vector Free Energy Calculations Calculating Free Energies Using Average Force. J. Chem. Phys. 2001, 115, 9169–9183.

(64) Feller, S. E.; Zhang, Y.; Pastor, R. W.; Brooks, B. R. Constant Pressure Molecular Dynamics Simulation: The Langevin Piston Method. J. Chem. Phys. 1995, 103 (11), 4613–4621.

(65) Lomize, A. L.; Pogozheva, I. D.; Mosberg, H. I. Anisotropic Solvent Model of the Lipid Bilayer. 2. Energetics of Insertion of Small Molecules, Peptides, and Proteins in Membranes. J. Chem. Inf. Model. 2011, 51 (4), 930–946.

(66) Tack, B. F.; Sawai, M. V; Kearney, W. R.; Robertson, A. D.; Sherman, M. A.; Wang, W.; Hong, T.; Boo, L. M.; Wu, H.; Waring, A. J.; Lehrer, R. I. SMAP-29 Has Two LPS-Binding Sites and a Central Hinge. Eur. J. Biochem. 2002, 269 (4), 1181–1189.

(67) Szolajska, E.; Poznanski, J.; Ferber, M. L.; Michalik, J.; Gout, E.; Fender, P.; Bailly, I.; Dublet, B.; Chroboczek, J. Poneratoxin, a Neurotoxin from Ant Venom: Structure and Expression in Insect Cells and Construction of a Bio-Insecticide. Eur. J. Biochem. 2004, 271 (11), 2127–2136.

(68) Strandberg, E.; Esteban-Martín, S.; Salgado, J.; Ulrich, A. S. Orientation and Dynamics of Peptides in Membranes Calculated from 2H-NMR Data. Biophys. J. 2009, 96 (8), 3223–3232.

(69) Yano, Y.; Matsuzaki, K. Measurement of Thermodynamic Parameters for Hydrophobic Mismatch 1: Self-Association of a Transmembrane Helix. Biochemistry 2006, 45 (10), 3370–3378.

(70) Kim, T.; Im, W. Revisiting Hydrophobic Mismatch with Free Energy Simulation Studies of Transmembrane Helix Tilt and Rotation. Biophys. J. 2010, 99 (1), 175–183.

(71) Özdirekcan, S.; Etchebest, C.; Killian, J. A.; Fuchs, P. F. J. On the Orientation of a Designed Transmembrane Peptide: Toward the Right Tilt Angle? J. Am. Chem. Soc. 2007, 129 (49), 15174–15181.

(72) Monticelli, L.; Tieleman, D. P.; Fuchs, P. F. J. Interpretation Of2H-NMR Experiments on the Orientation of the Transmembrane Helix WALP23 by Computer Simulations. Biophys. J. 2010, 99 (5), 1455–1464.

(73) Fiorin, G.; Klein, M. L.; Henin, J. Using Collective Variables to Drive Molecular Dynamics Simulations. Mol. Phys. 2013, 111 (22–23), 3345–3362.

(74) Fisher, L. E.; Engelman, D. M.; Sturgis, J. N. Effect of Detergents on the Association of the Glycophorin A Transmembrane Helix. Biophys. J. 2003, 85 (5), 3097–3105.

(75) Fleming, K. G. Standardizing the Free Energy Change of Transmembrane Helix-Helix Interactions. J. Mol. Biol. 2002, 323 (3), 563–571.

(76) Chen, L.; Novicky, L.; Merzlyakov, M.; Hristov, T.; Hristova, K. Measuring the Energetics of Membrane Protein Dimerization in Mammalian Membranes. J. Am. Chem. Soc. 2010, 132 (10), 3628–3635.

(77) Sarabipour, S.; Hristova, K. Glycophorin A Transmembrane Domain Dimerization in Plasma Membrane Vesicles Derived from CHO, HEK 293T, and A431 Cells. Biochim. Biophys. Acta - Biomembr. 2013, 1828 (8), 1829–1833.

(78) Hénin, J.; Pohorille, A.; Chipot, C. Insights into the Recognition and Association of Transmembrane α-Helices. The Free Energy of α-Helix Dimerization in Glycophorin A. J. Am. Chem. Soc. 2005, 127 (23), 8478–8484.

(79) Sengupta, D.; Marrink, S. J. Lipid-Mediated Interactions Tune the Association of Glycophorin A Helix and Its Disruptive Mutants in Membranes. Phys. Chem. Chem. Phys. 2010, 12 (40), 12987–12996.

(80) North, B.; Cristian, L.; Fu Stowell, X.; Lear, J. D.; Saven, J. G.; DeGrado, W. F. Characterization of a Membrane Protein Folding Motif, the Ser Zipper, Using Designed Peptides. J. Mol. Biol. 2006, 359 (4), 930–939.

(81) Castillo, N.; Monticelli, L.; Barnoud, J.; Tieleman, D. P. Free Energy of WALP23 Dimer Association in DMPC, DPPC, and DOPC Bilayers. Chem. Phys. Lipids 2013, 169, 95–105.

(82) Artemenko, E. O.; Egorova, N. S.; Arseniev, A. S.; Feofanov, A. V. Transmembrane Domain of EphA1 Receptor Forms Dimers in Membrane-like Environment. Biochim. Biophys. Acta - Biomembr. 2008, 1778 (10), 2361–2367.

(83) Chavent, M.; Chetwynd, A. P.; Stansfeld, P. J.; Sansom, M. S. P. Dimerization of the EphA1 Receptor Tyrosine Kinase Transmembrane Domain: Insights into the Mechanism of Receptor Activation. Biochemistry 2014, 53 (42), 6641–6652.

(84) Javanainen, M.; Martinez-Seara, H.; Vattulainen, I. Excessive Aggregation of Membrane Proteins in the Martini Model. PLoS One 2017, 12 (11), e0187936.

(85) Chen, L.; Merzlyakov, M.; Cohen, T.; Shai, Y.; Hristova, K. Energetics of ErbB1 Transmembrane Domain Dimerization in Lipid Bilayers. Biophys. J. 2009, 96 (11), 4622–4630.

(86) Lelimousin, M.; Limongelli, V.; Sansom, M. S. P. Conformational Changes in the Epidermal Growth Factor Receptor: Role of the Transmembrane Domain Investigated by Coarse-Grained MetaDynamics Free Energy Calculations. J. Am. Chem. Soc. 2016, 138 (33), 10611–10622.

(87) Simons, K. T.; Kooperberg, C.; Huang, E.; Baker, D. Assembly of Protein Tertiary Structures from Fragments with Similar Local Sequences Using Simulated Annealing and Bayesian Scoring Functions. J. Mol. Biol. 1997, 268 (1), 209–225.

(88) Samudrala, R.; Levitt, M. Decoys ‘R’ Us: A Database of Incorrect Conformations to Improve Protein Structure Prediction.’ Protein Sci. 2000, 9 (7), 1399–1401.

(89) Simons, K. T.; Ruczinski, I.; Kooperberg, C.; Fox, B. A.; Bystroff, C.; Baker, D. Improved Recognition of Native-like Protein Structures Using a Combination of Sequence-Dependent and Sequence-Independent Features of Proteins. Proteins Struct. Funct. Genet. 1999, 34 (1), 82–95.

(90) Keasar, C.; Levitt, M. A Novel Approach to Decoy Set Generation: Designing a Physical Energy Function Having Local Minima with Native Structure Characteristics. J. Mol. Biol. 2003, 329 (1), 159–174.

(91) Xia, Y.; Huang, E. S.; Levitt, M.; Samudrala, R. Ab Initio Construction of Protein Tertiary Structures Using a Hierarchical Approach. J. Mol. Biol. 2000, 300 (1), 171– 185.

(92) Lee, S. Y.; Lee, A.; Chen, J.; MacKinnon, R. Structure of the KvAP Voltage-Dependent K+ Channel and Its on the Lipid Membrane. Proc. Natl. Acad. Sci. U. S. A. 2005, 102 (43), 15441–15446.

(93) Liu, Z.; Gandhi, C. S.; Rees, D. C. Structure of a Tetrameric MscL in an Expanded Intermediate State. Nature 2009, 461 (7260), 120–124.

(94) Bevers, E. M.; Williamson, P. L. Phospholipid Scramblase: An Update. Febs Lett. 2010, 584 (13), 2724–2730.

(95) Bethel, N. P.; Grabe, M. Atomistic Insight into Lipid Translocation by a TMEM16 Scramblase. Proc. Natl. Acad. Sci. 2016, 113 (49), 14049–14054.

(96) Jiang, T.; Yu, K.; Hartzell, H. C.; Tajkhorshid, E. Lipids and Ions Traverse the Membrane by the Same Physical Pathway in the NhTMEM16 Scramblase. Elife 2017, 6, e28671.

(97) Lee, B.-C.; Khelashvili, G.; Falzone, M.; Menon, A. K.; Weinstein, H.; Accardi, A. Gating Mechanism of the Extracellular Entry to the Lipid Pathway in a TMEM16 Scramblase. Nat. Commun. 2018, 9 (1), 3251.

(98) Andoh, Y.; Yoshii, N.; Yamada, A.; Fujimoto, K.; Kojima, H.; Mizutani, K.; Nakagawa, A.; Nomoto, A.; Okazaki, S. All-Atom Molecular Dynamics Calculation Study of Entire Poliovirus Empty Capsids in Solution. J. Chem. Phys. 2014, 141 (16), 165101.

(99) Zoni, V.; Khaddaj, R.; Lukmantara, I.; Shinoda, W.; Yang, H.; Schneiter, R.; Vanni, S. Seipin Accumulates and Traps Diacylglycerols and Triglycerides in Its Ring-like Structure. Proc. Natl. Acad. Sci. 2021, 118 (10), e2017205118.

