## Supplemental Figures for "The SPICA Coarse-Grained Force Field for Proteins and Peptides"

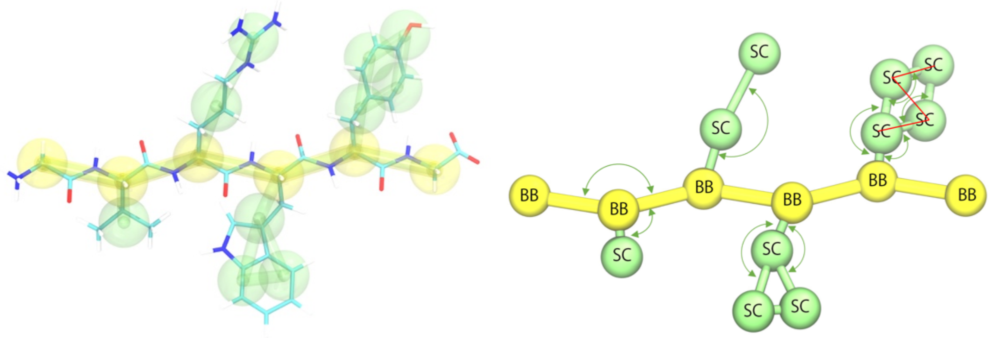


Figure S1. Schematic representation of the five different geometrical classes of amino acids, consisting of either zero, one, two, three, or four beads for the side chain (plus a backbone bead). The yellow lines are bond interaction, blue curves are angle interaction, and red lines are dihedral interaction. Backbone beads are indicated by “B” and side-chain beads by “S”. Harmonic angle potential is added on all B-B-B, B-B-S and B-S-S angles and two of four S-S-S angles in four beads side chain. Dihedral potential is only used for four beads side chain of PHE and TYR.

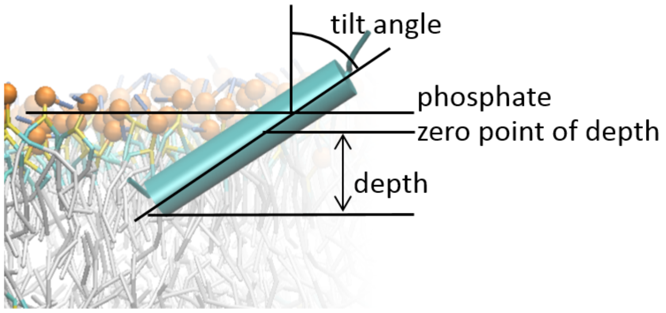


Figure S2. A schematic diagram for the calculation of penetration depth and tilt angle of a peptide/protein.

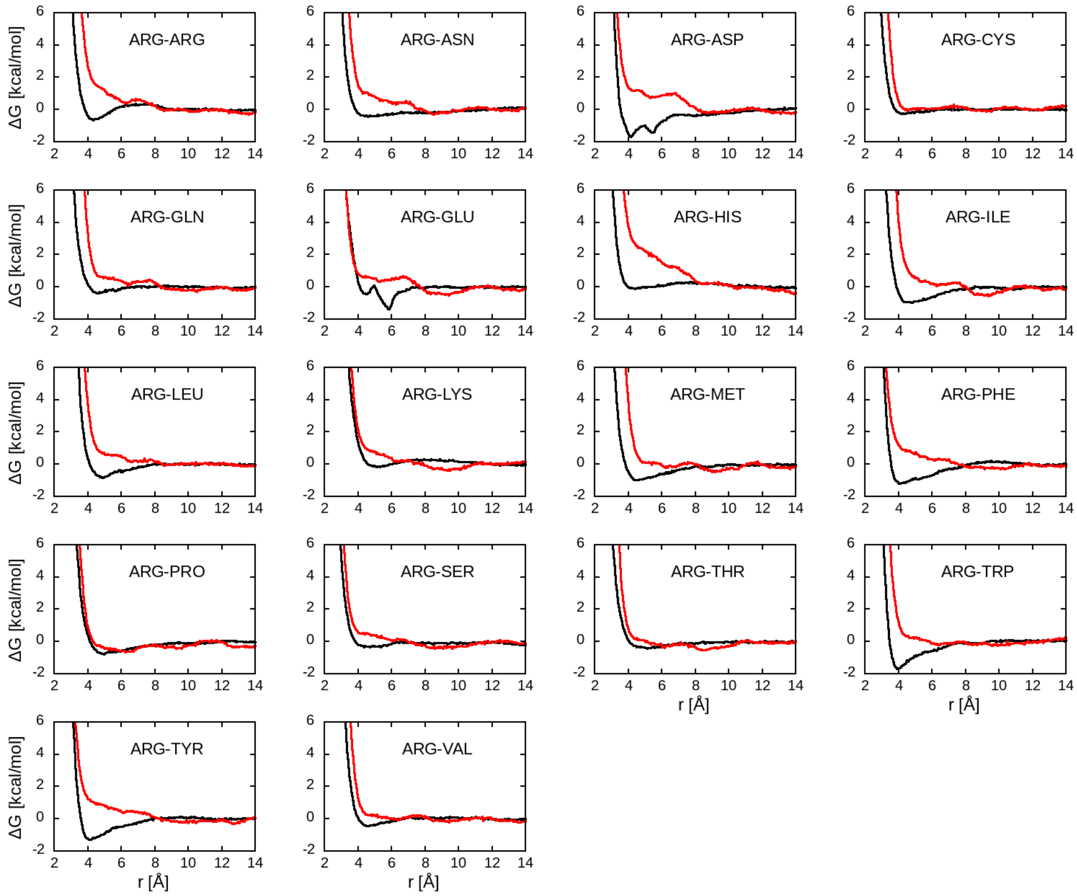
ΔG [kcal/mol]
r [Å]
ARG-ARG
ARG-ASN
ARG-ASP
ARG-CYS
ARG-GLN
ARG-GLU
ARG-HIS
ARG-ILE
ARG-LEU
ARG-LYS
ARG-MET
ARG-PHE
ARG-PRO
ARG-SER
ARG-THR
ARG-TRP
ARG-TYR
ARG-VAL

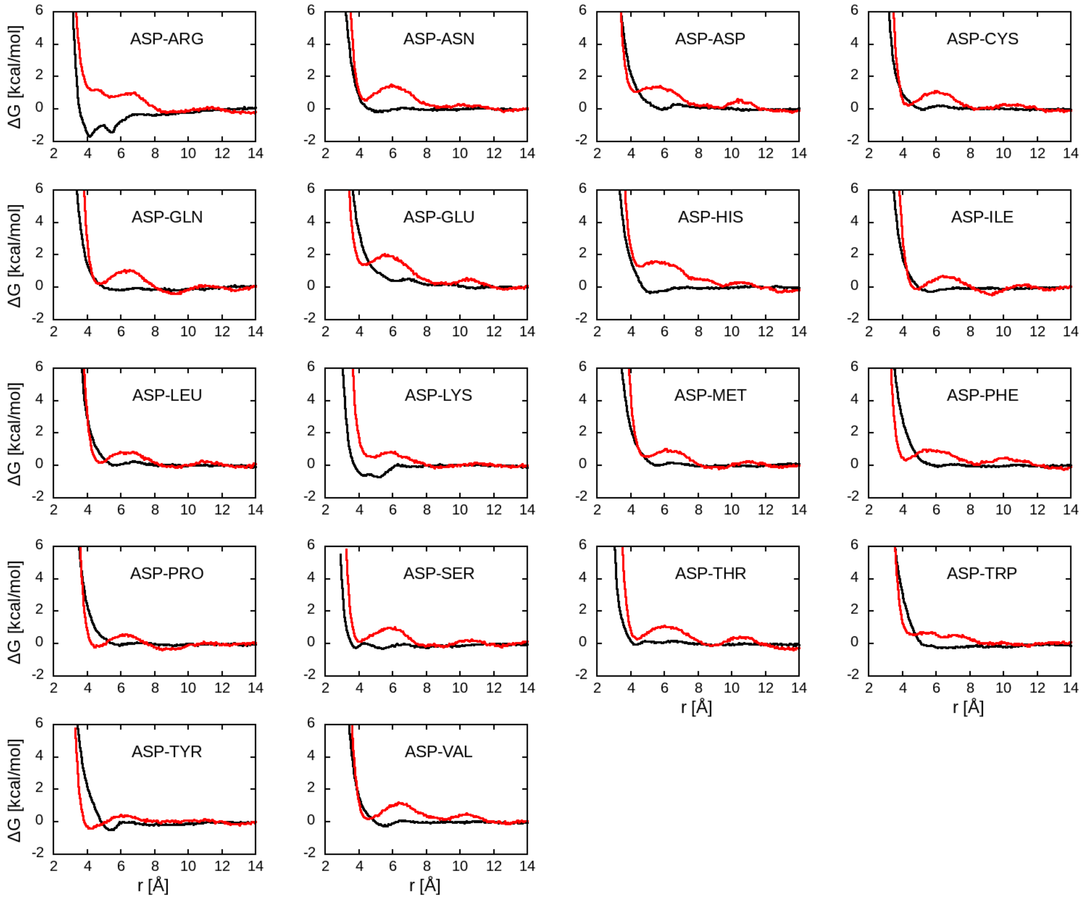
ΔG [kcal/mol]
r [Å]
ASP-ARG
ASP-ASN
ASP-ASP
ASP-CYS
ASP-GLN
ASP-GLU
ASP-HIS
ASP-ILE
ASP-LEU
ASP-LYS
ASP-MET
ASP-PHE
ASP-PRO
ASP-SER
ASP-THR
ASP-TRP
ASP-TYR
ASP-VAL

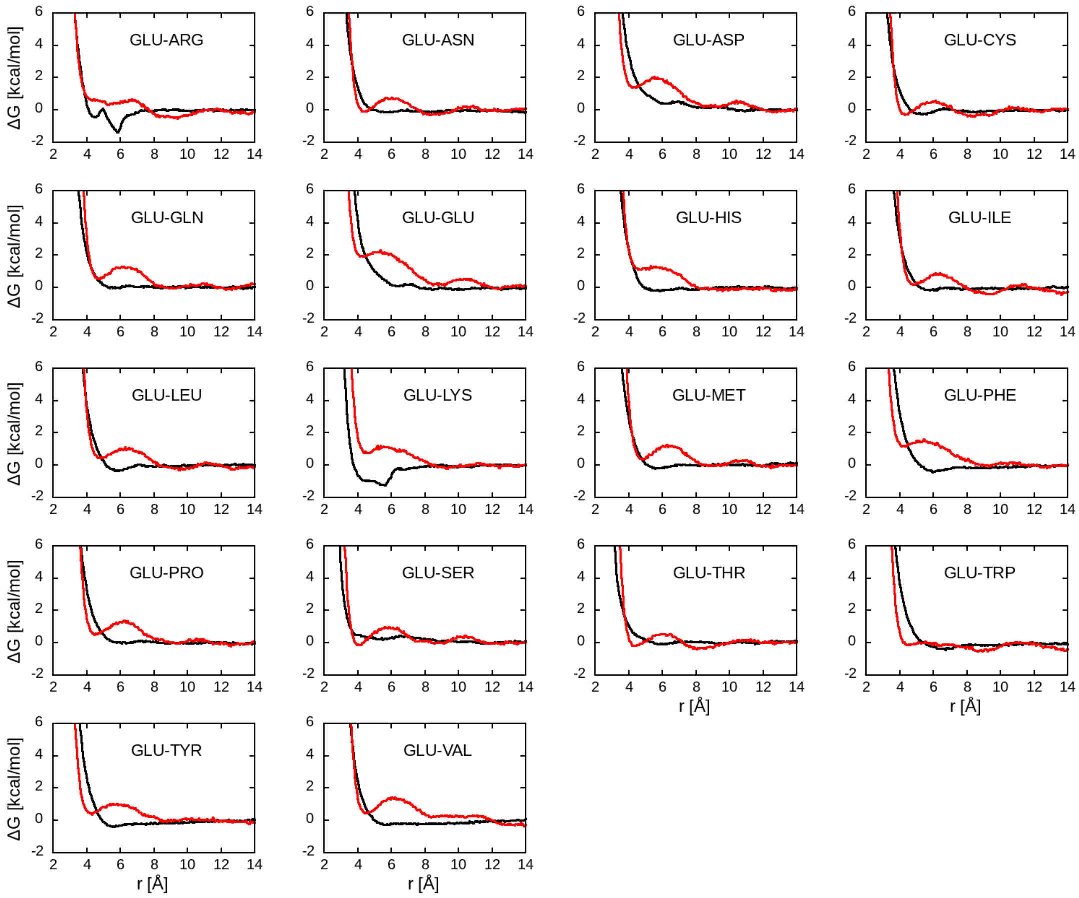
GLU-ARG
GLU-ASN
GLU-ASP
GLU-CYS
GLU-GLN
GLU-GLU
GLU-HIS
GLU-ILE
GLU-LEU
GLU-LYS
GLU-MET
GLU-PHE
GLU-PRO
GLU-SER
GLU-THR
GLU-TRP
GLU-TYR
GLU-VAL
ΔG [kcal/mol]
r [Å]

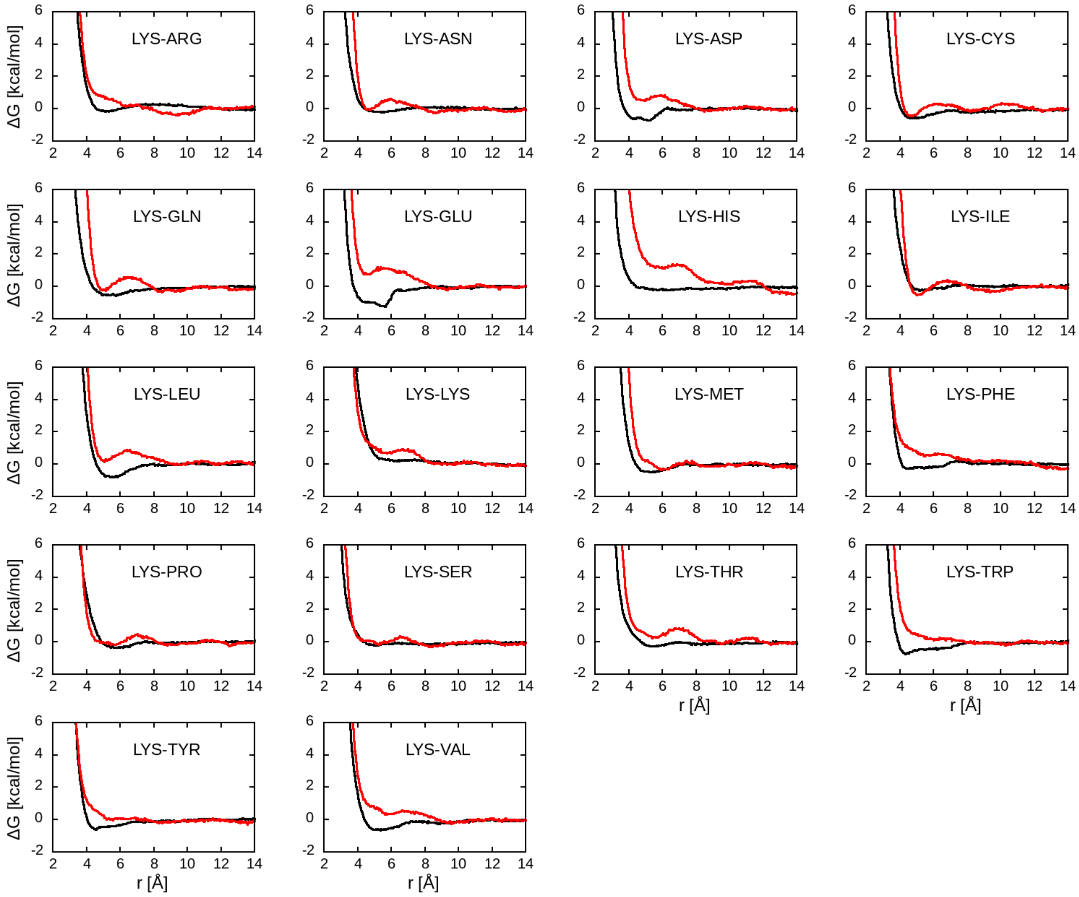
LYS-ARG
LYS-ASN
LYS-ASP
LYS-CYS
LYS-GLN
LYS-GLU
LYS-HIS
LYS-ILE
LYS-LEU
LYS-LYS
LYS-MET
LYS-PHE
LYS-PRO
LYS-SER
LYS-THR
LYS-TRP
LYS-TYR
LYS-VAL
ΔG [kcal/mol]
r [Å]

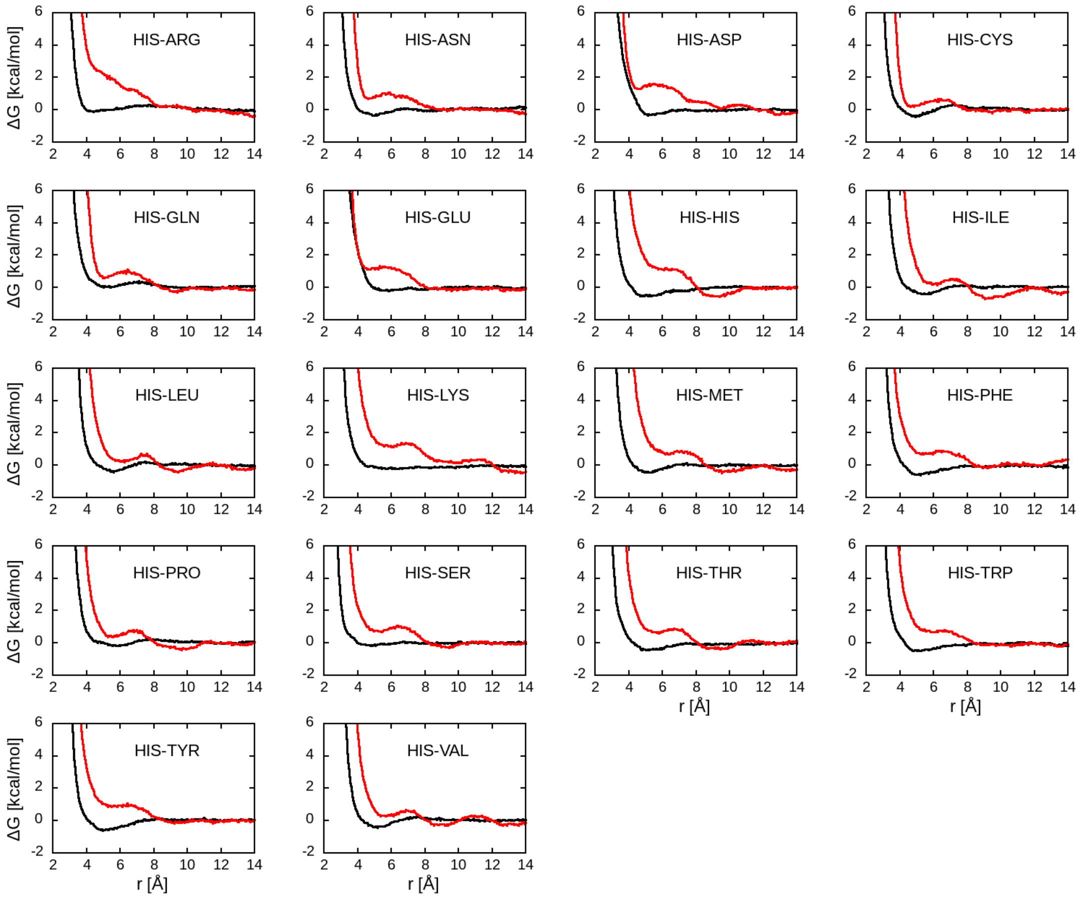


Figure S3. Dimerization free energies in kcal/mol of side chain analogues for charged and HIS residues as a function of distance in Å. Black lines are calculated using CHARMM all-atom force field, and red lines are calculated using the initial LJ parameters employing the combination rules.

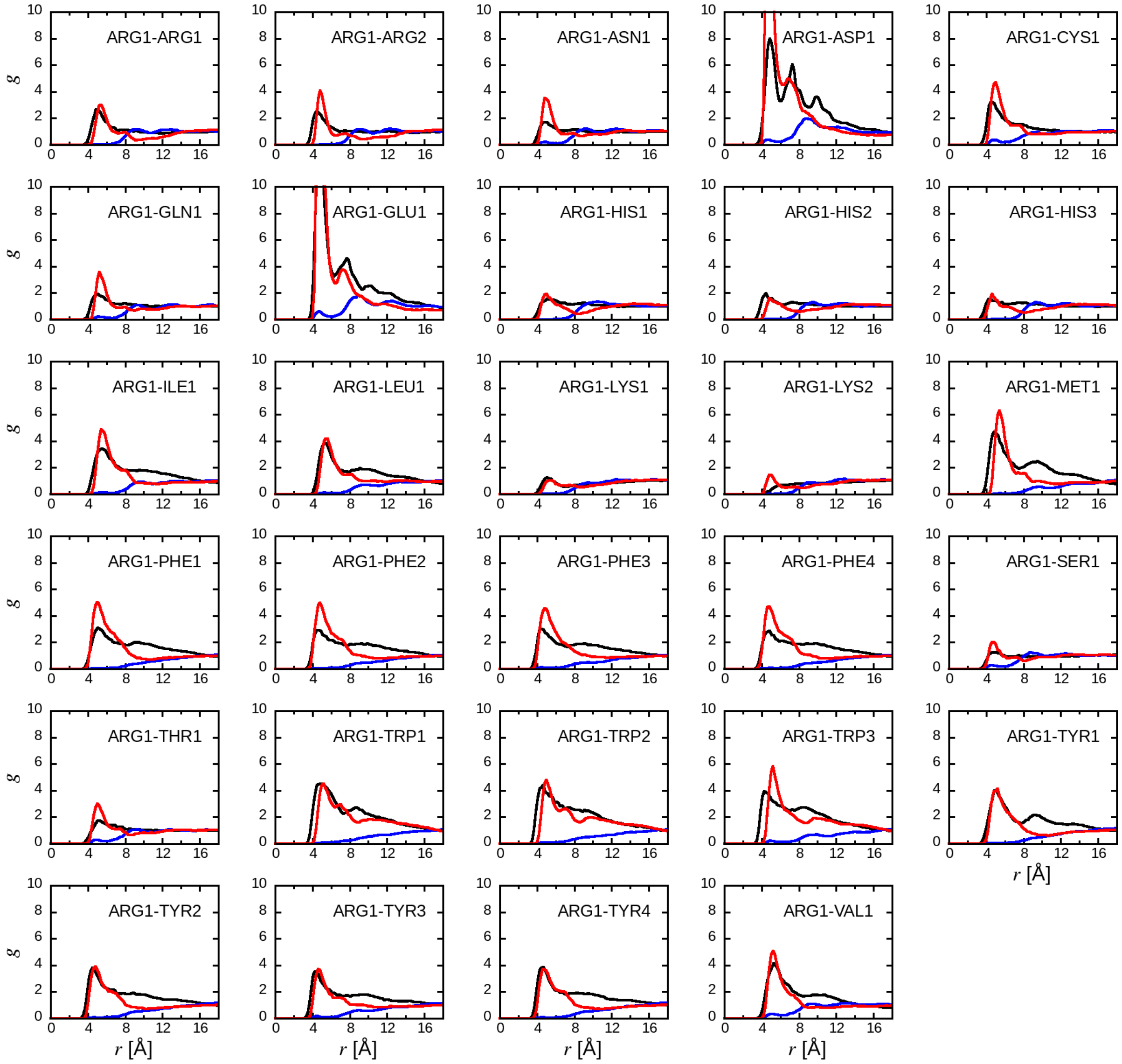

ARG1-ARG1
ARG1-ARG2
ARG1-ASN1
ARG1-ASP1
ARG1-CYS1
ARG1-GLN1
ARG1-GLU1
ARG1-HIS1
ARG1-HIS2
ARG1-HIS3
ARG1-ILE1
ARG1-LEU1
ARG1-LYS1
ARG1-LYS2
ARG1-MET1
ARG1-PHE1
ARG1-PHE2
ARG1-PHE3
ARG1-PHE4
ARG1-SER1
ARG1-THR1
ARG1-TRP1
ARG1-TRP2
ARG1-TRP3
ARG1-TYR1
ARG1-TYR2
ARG1-TYR3
ARG1-TYR4
ARG1-VAL1
g
r [Å]

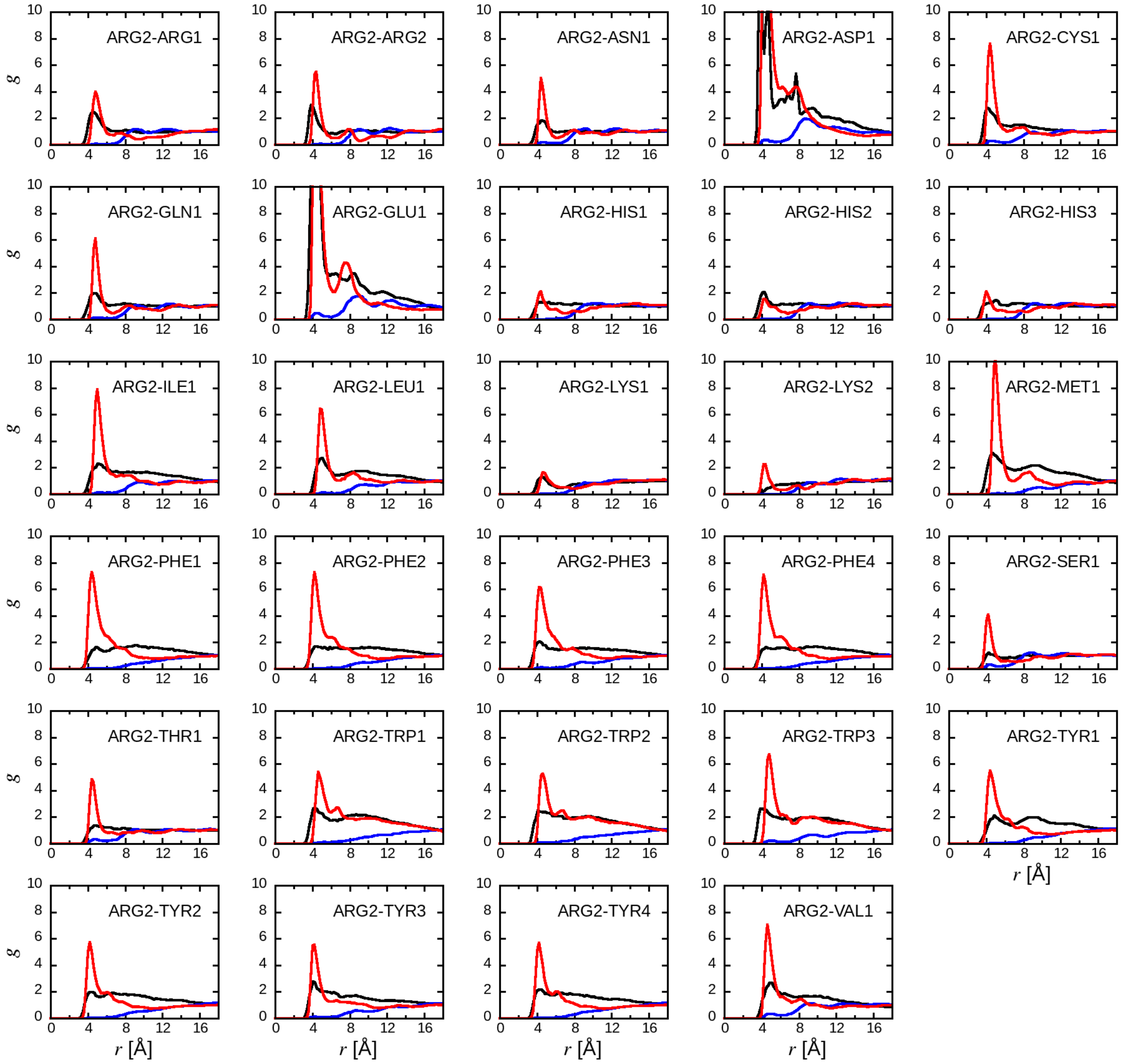
ARG2-ARG1
ARG2-ARG2
ARG2-ASN1
ARG2-ASP1
ARG2-CYS1
ARG2-GLN1
ARG2-GLU1
ARG2-HIS1
ARG2-HIS2
ARG2-HIS3
ARG2-ILE1
ARG2-LEU1
ARG2-LYS1
ARG2-LYS2
ARG2-MET1
ARG2-PHE1
ARG2-PHE2
ARG2-PHE3
ARG2-PHE4
ARG2-SER1
ARG2-THR1
ARG2-TRP1
ARG2-TRP2
ARG2-TRP3
ARG2-TYR1
ARG2-TYR2
ARG2-TYR3
ARG2-TYR4
ARG2-VAL1
g
r [Å]

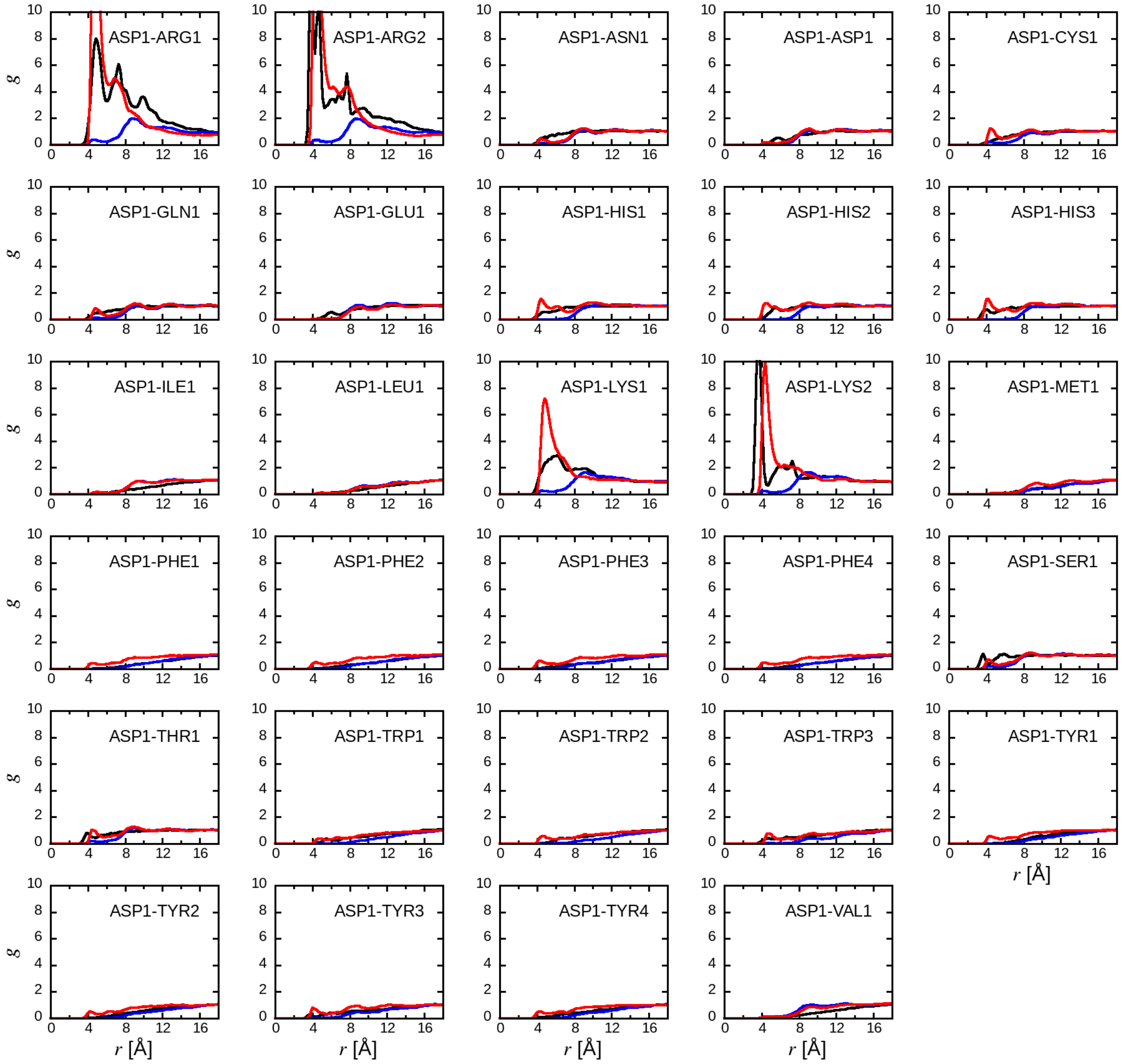
ASP1-ARG1
ASP1-ARG2
ASP1-ASN1
ASP1-ASP1
ASP1-CYS1
ASP1-GLN1
ASP1-GLU1
ASP1-HIS1
ASP1-HIS2
ASP1-HIS3
ASP1-ILE1
ASP1-LEU1
ASP1-LYS1
ASP1-LYS2
ASP1-MET1
ASP1-PHE1
ASP1-PHE2
ASP1-PHE3
ASP1-PHE4
ASP1-SER1
ASP1-THR1
ASP1-TRP1
ASP1-TRP2
ASP1-TRP3
ASP1-TYR1
ASP1-TYR2
ASP1-TYR3
ASP1-TYR4
ASP1-VAL1
g
r [Å]

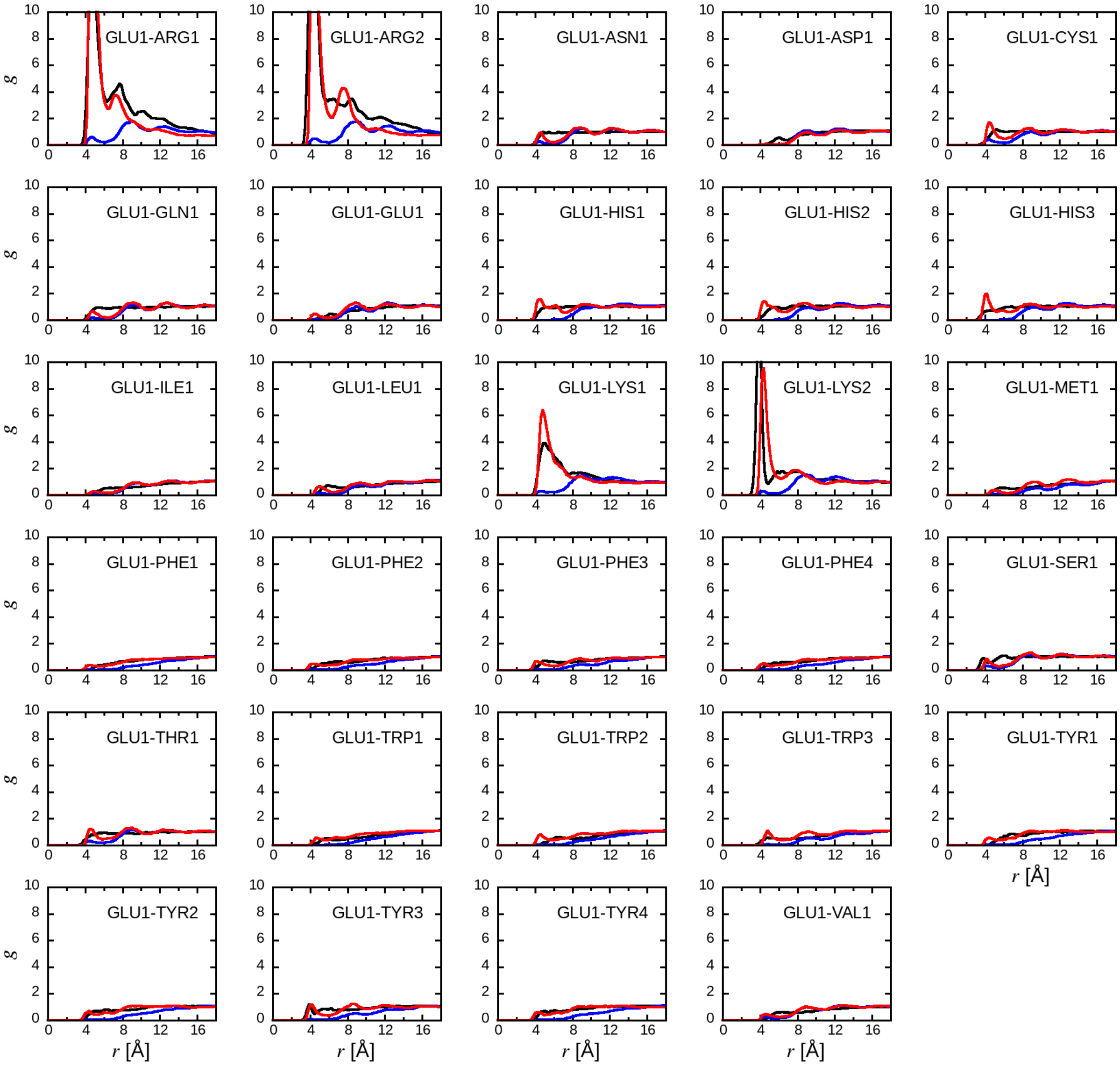
GLU1-ARG1
GLU1-ARG2
GLU1-ASN1
GLU1-ASP1
GLU1-CYS1
GLU1-GLN1
GLU1-GLU1
GLU1-HIS1
GLU1-HIS2
GLU1-HIS3
GLU1-ILE1
GLU1-LEU1
GLU1-LYS1
GLU1-LYS2
GLU1-MET1
GLU1-PHE1
GLU1-PHE2
GLU1-PHE3
GLU1-PHE4
GLU1-SER1
GLU1-THR1
GLU1-TRP1
GLU1-TRP2
GLU1-TRP3
GLU1-TYR1
GLU1-TYR2
GLU1-TYR3
GLU1-TYR4
GLU1-VAL1
g
r [Å]

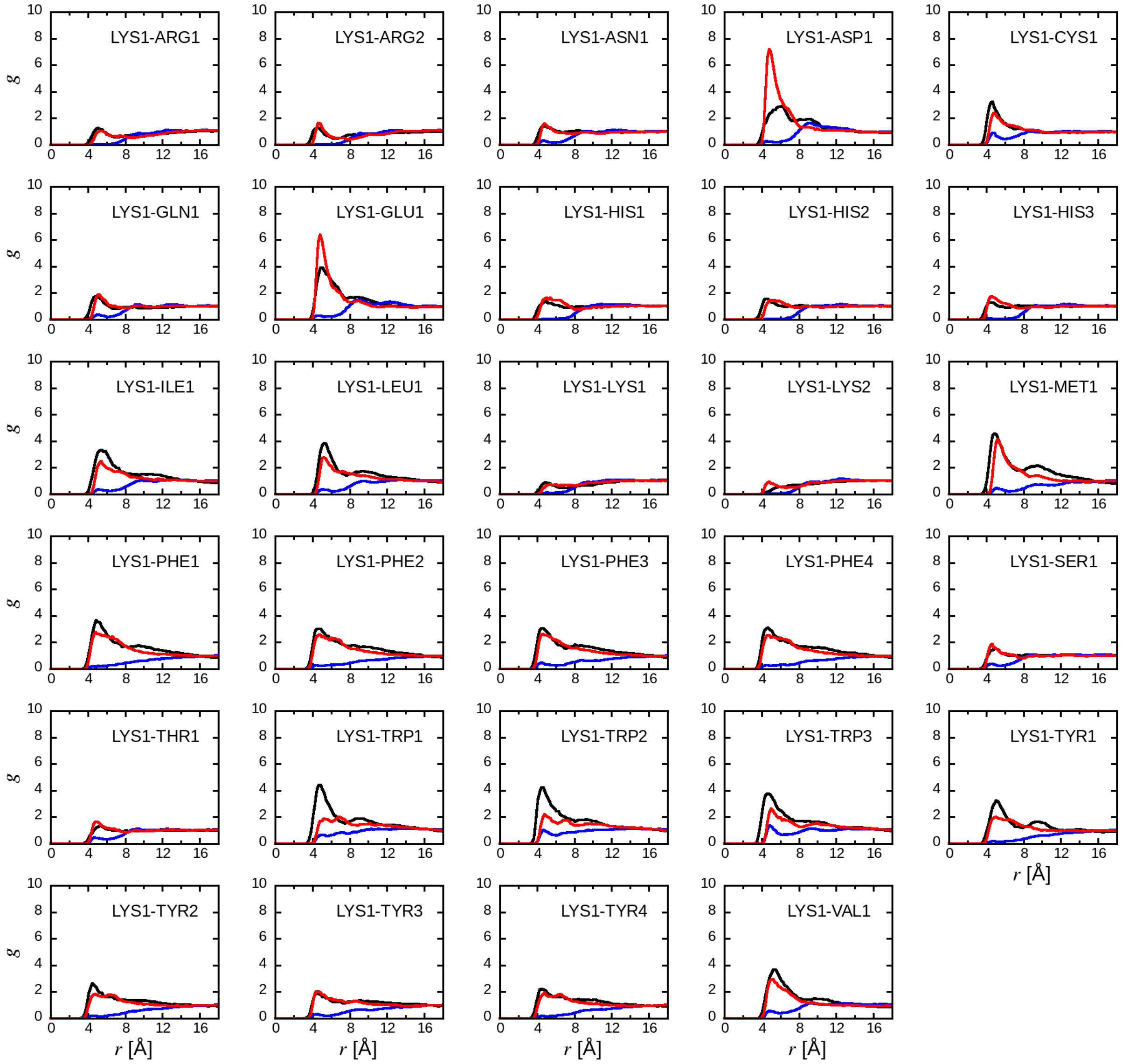

LYS1-ARG1
LYS1-ARG2
LYS1-ASN1
LYS1-ASP1
LYS1-CYS1
LYS1-GLN1
LYS1-GLU1
LYS1-HIS1
LYS1-HIS2
LYS1-HIS3
LYS1-ILE1
LYS1-LEU1
LYS1-LYS1
LYS1-LYS2
LYS1-MET1
LYS1-PHE1
LYS1-PHE2
LYS1-PHE3
LYS1-PHE4
LYS1-SER1
LYS1-THR1
LYS1-TRP1
LYS1-TRP2
LYS1-TRP3
LYS1-TYR1
LYS1-TYR2
LYS1-TYR3
LYS1-TYR4
LYS1-VAL1
g
r [Å]

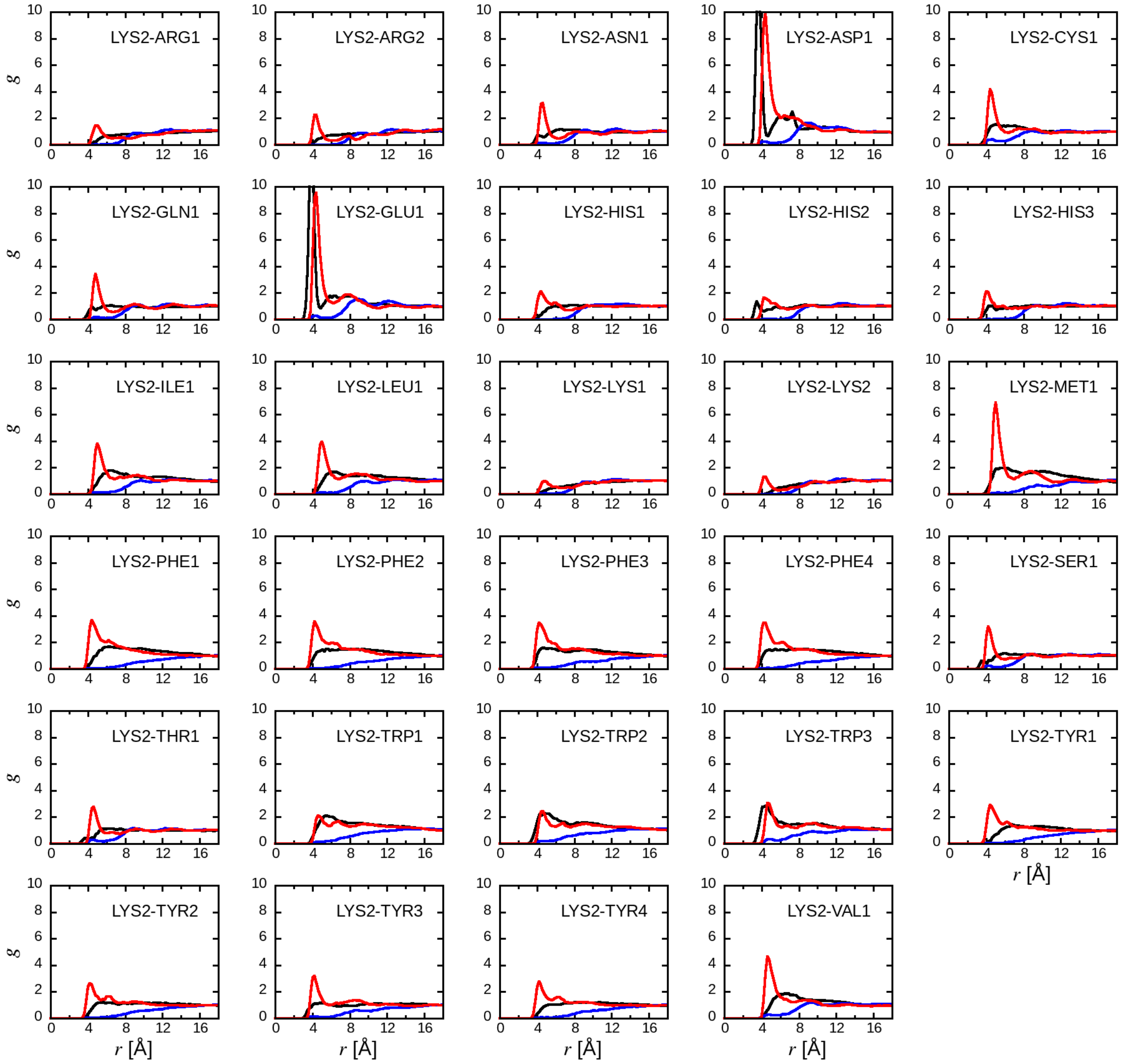
LYS2-ARG1
LYS2-ARG2
LYS2-ASN1
LYS2-ASP1
LYS2-CYS1
LYS2-GLN1
LYS2-GLU1
LYS2-HIS1
LYS2-HIS2
LYS2-HIS3
LYS2-ILE1
LYS2-LEU1
LYS2-LYS1
LYS2-LYS2
LYS2-MET1
LYS2-PHE1
LYS2-PHE2
LYS2-PHE3
LYS2-PHE4
LYS2-SER1
LYS2-THR1
LYS2-TRP1
LYS2-TRP2
LYS2-TRP3
LYS2-TYR1
LYS2-TYR2
LYS2-TYR3
LYS2-TYR4
LYS2-VAL1
g
r [Å]

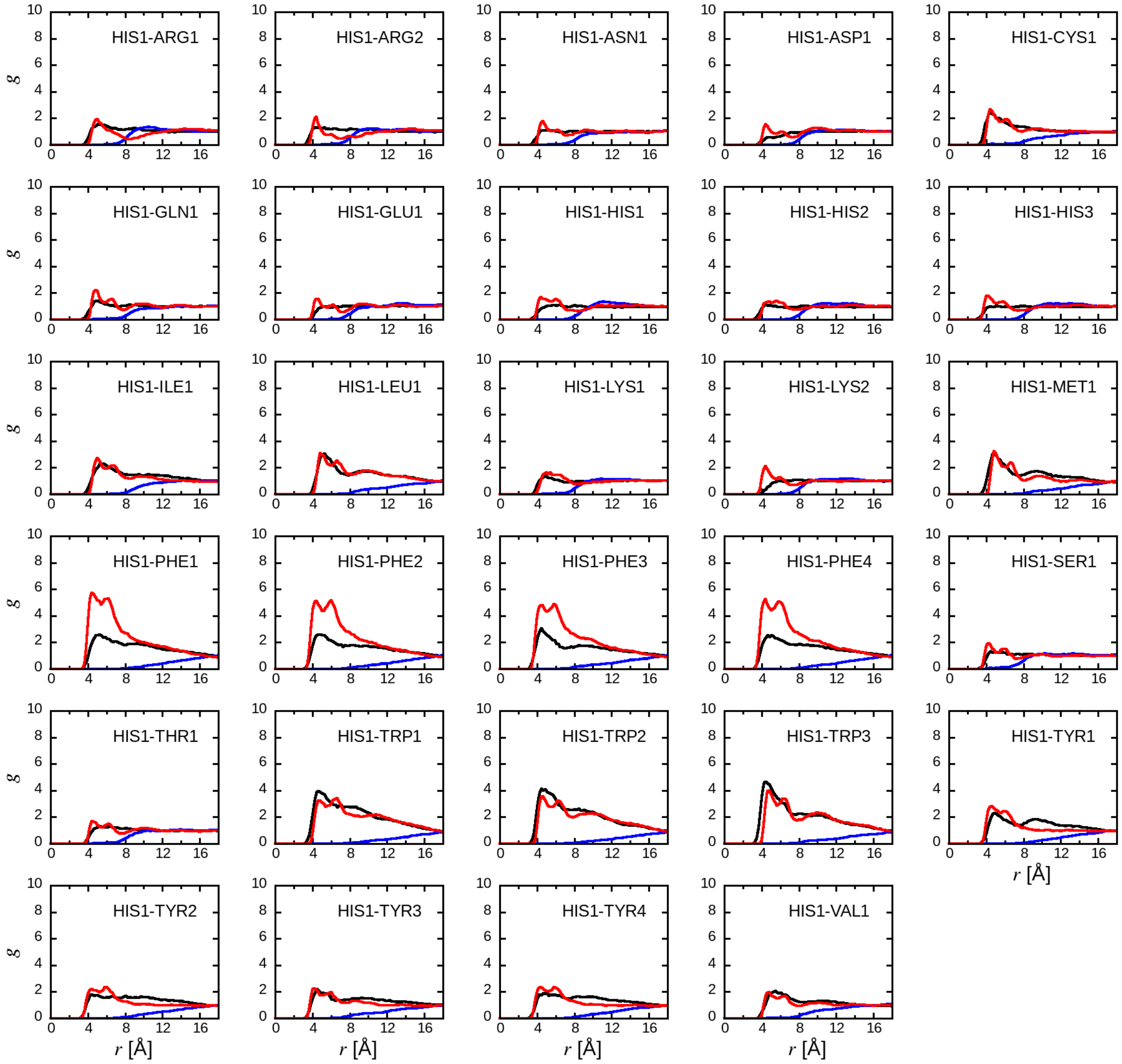

HIS1-ARG1
HIS1-ARG2
HIS1-ASN1
HIS1-ASP1
HIS1-CYS1
HIS1-GLN1
HIS1-GLU1
HIS1-HIS1
HIS1-HIS2
HIS1-HIS3
HIS1-ILE1
HIS1-LEU1
HIS1-LYS1
HIS1-LYS2
HIS1-MET1
HIS1-PHE1
HIS1-PHE2
HIS1-PHE3
HIS1-PHE4
HIS1-SER1
HIS1-THR1
HIS1-TRP1
HIS1-TRP2
HIS1-TRP3
HIS1-TYR1
HIS1-TYR2
HIS1-TYR3
HIS1-TYR4
HIS1-VAL1
g
r [Å]

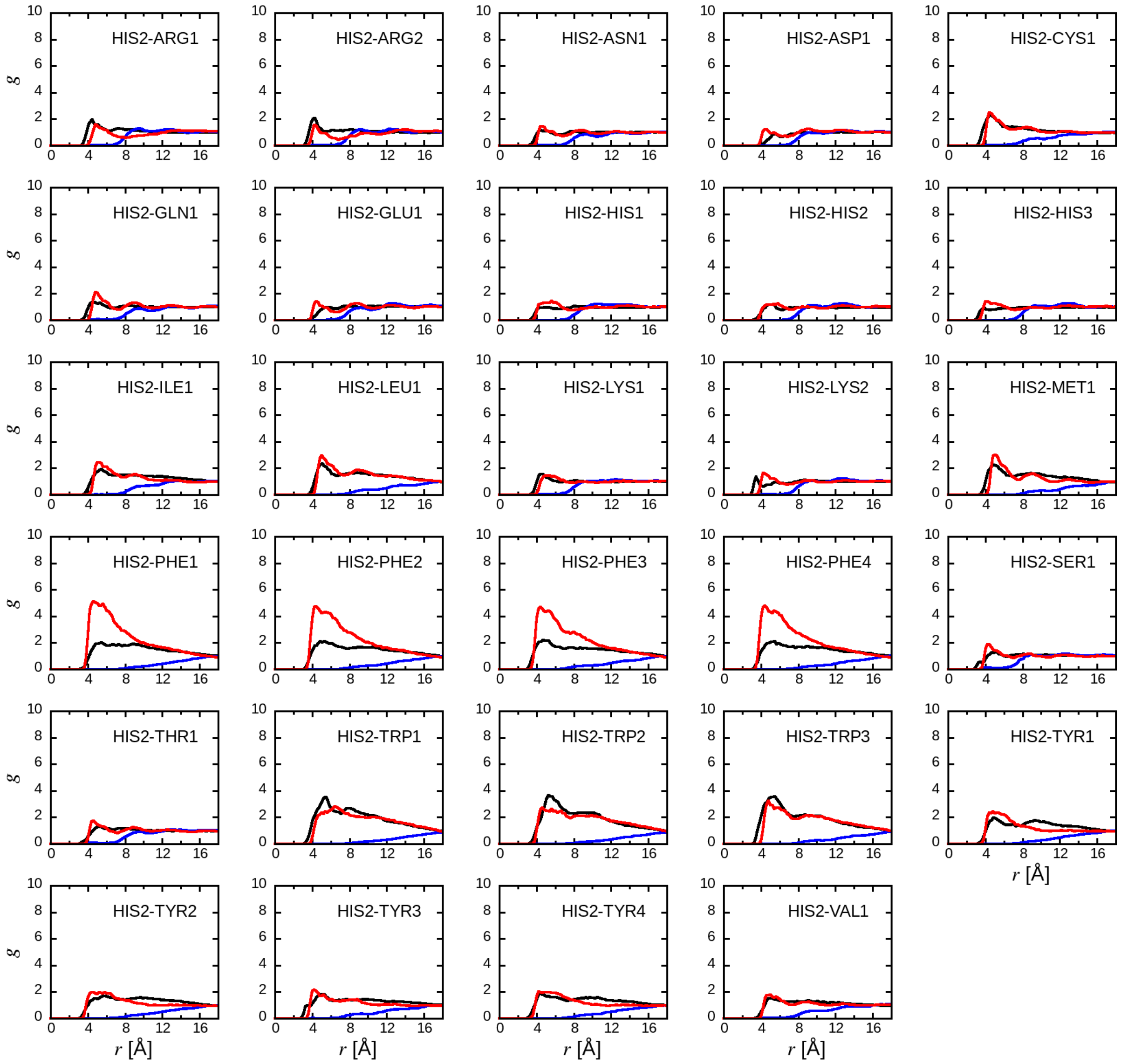
HIS2-ARG1
HIS2-ARG2
HIS2-ASN1
HIS2-ASP1
HIS2-CYS1
HIS2-GLN1
HIS2-GLU1
HIS2-HIS1
HIS2-HIS2
HIS2-HIS3
HIS2-ILE1
HIS2-LEU1
HIS2-LYS1
HIS2-LYS2
HIS2-MET1
HIS2-PHE1
HIS2-PHE2
HIS2-PHE3
HIS2-PHE4
HIS2-SER1
HIS2-THR1
HIS2-TRP1
HIS2-TRP2
HIS2-TRP3
HIS2-TYR1
HIS2-TYR2
HIS2-TYR3
HIS2-TYR4
HIS2-VAL1
g
r [Å]

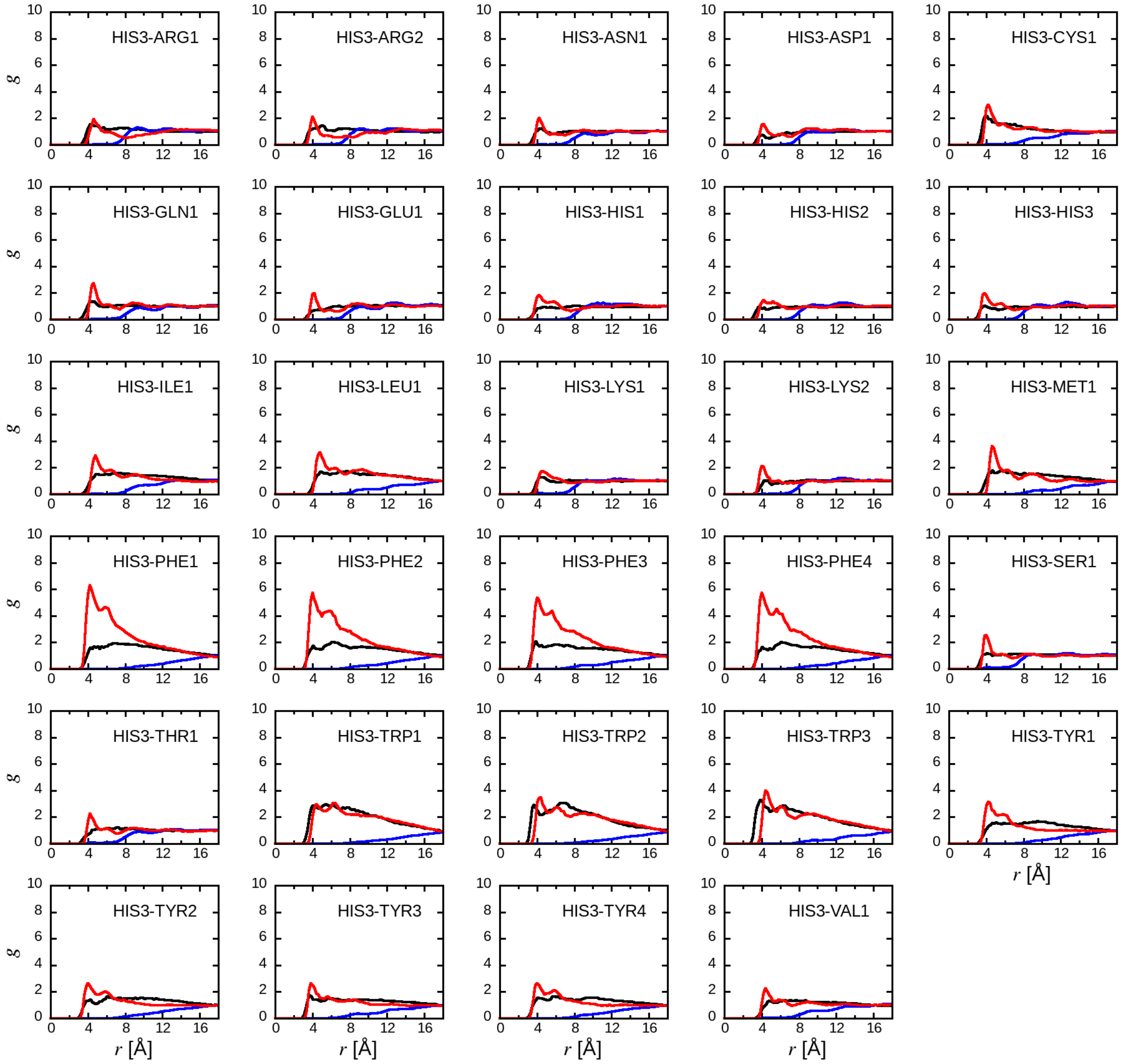


Figure S4. The radial distribution function, $g$, between amino acid side chain analogue dimers in aqueous solution. Particularly the amino acid side chain analogues of the charged and HIS segments are plotted. Black lines are the results from AA MD. Blue lines are the result of the CG MD using the LJ parameters guessed by applying the combination rules. Red lines represent the results using the optimized SPICA CG model.

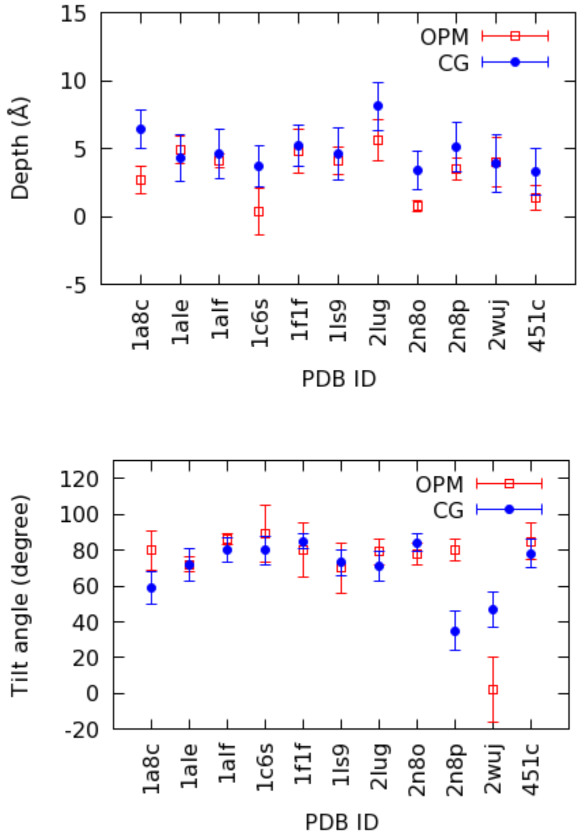


Figure S5. Penetration depths and tilt angles of peripheral proteins.

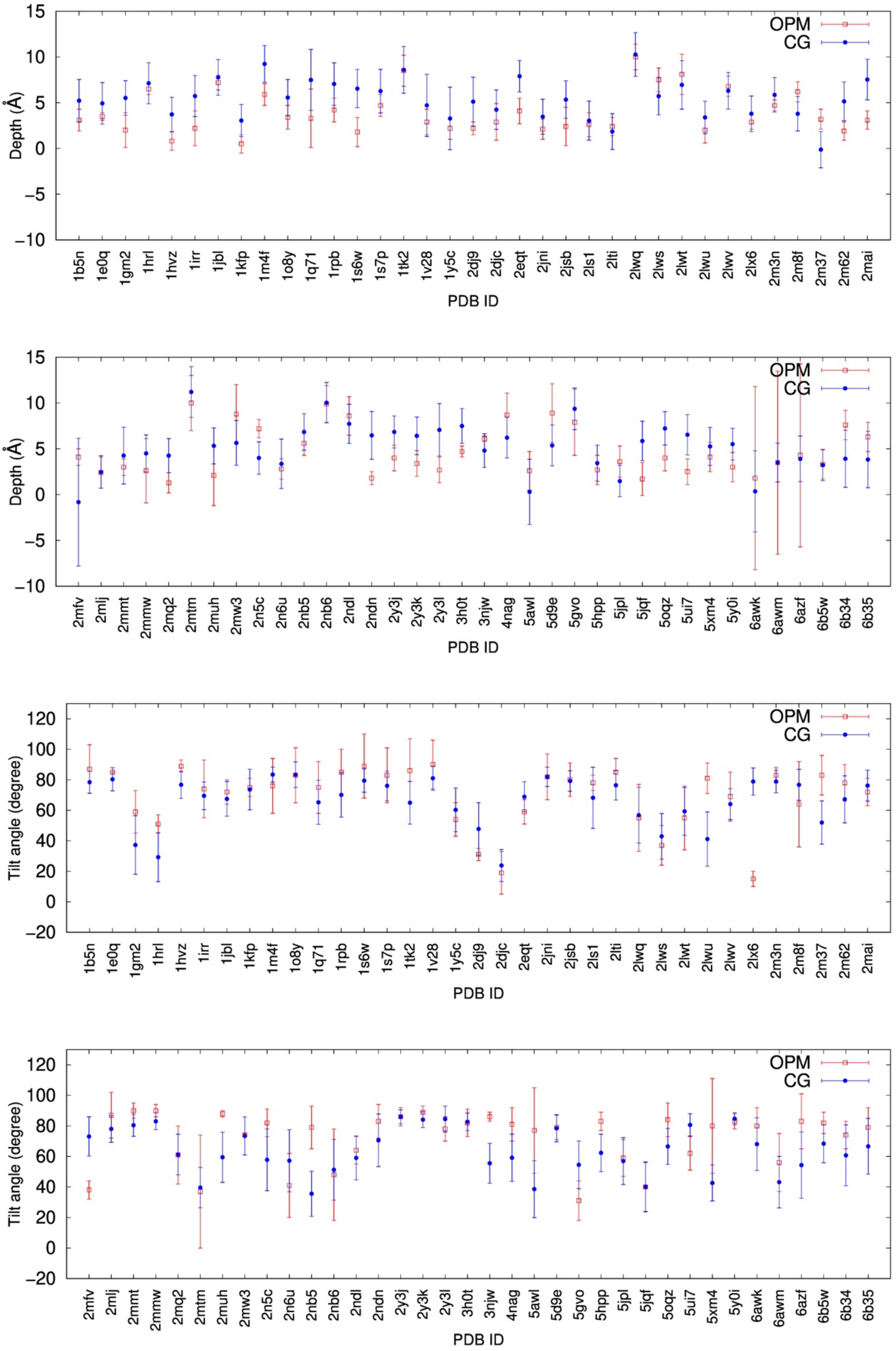


Figure S6: Penetration depths and tilt angles of peripheral beta-hairpin peptides to DOPC membranes.

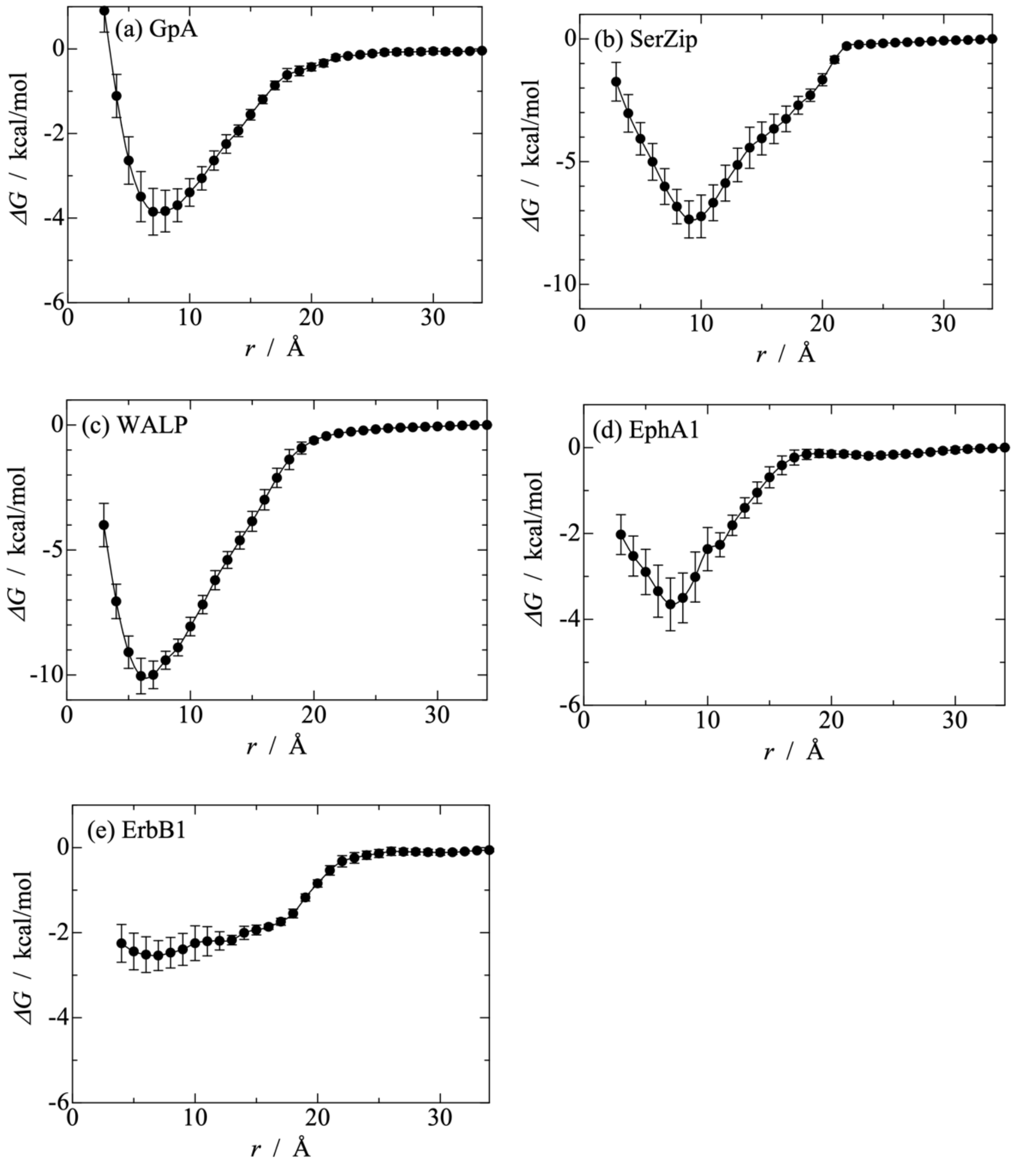


Figure S7. Dimerization free energy of transmembrane peptides. (a) gpAs in a DPPC bilayer at 323 K, (b) SerZips in a DMPC bilayer at 310 K, (c) WALPs in a DOPC bilayer at 308 K, (d) EphA1s in a DMPC bilayer at 303 K, and (e) ErbB1s in a DLPC bilayer at 303 K.
